# Lack of adaptation to centriolar defects leads to p53-independent microcephaly in the absence of Cep135

**DOI:** 10.1101/2020.05.07.082032

**Authors:** José González-Martínez, Andrzej W Cwetsch, Diego Martínez-Alonso, Luis Rodrigo López-Sainz, Jorge Almagro, Diego Megías, Jasminka Boskovic, Javier Gilabert-Juan, Osvaldo Graña-Castro, Alessandra Pierani, Axel Behrens, Sagrario Ortega, Marcos Malumbres

## Abstract

Autosomal Recessive Primary Microcephaly (MCPH) is a rare disease associated to proteins involved in centrosomal and spindle dynamics including Cep135 (MCPH8). Although Cep135 has been associated to centriolar assembly, the mechanisms associated to the pathogenesis underlying *MCPH8* mutations are unclear. By using a series of CRISPR/Cas9-edited murine *Cep135* alleles, we report here that lack of Cep135 results in perinatal lethality accompanied by significant microcephaly in a dosis-dependent manner. Cep135 deficiency, but not that of other centrosomal microcephaly proteins such as Aspm or Cdk5rap2, induces centrosome duplication defects, and perturbed centriole structure and dynamics. Whereas other cell types are able to quickly adapt to these defects, neural progenitors display a prolonged response leading to chromosomal instability and cell death in later developmental stages. Genetic ablation of *Trp53* in these mutant embryos prevents apoptotic cell death but does not rescue the microcephaly induced by Cep135 loss. These results suggest that microcephaly can arise from the lack of adaptation to centriole defects in neural progenitors of the developing neocortex in a p53-independent manner.

## Introduction

Autosomal Recessive Primary Microcephaly (MCPH) is a congenital brain disorder characterized by a reduction in head circumference linked to a striking decrease in brain volume (from −3 to −13 standard deviations). These changes are not linked with gross anomalies of brain architecture and associate with a primary and selective defect in the production of neurons during development. Several genes mutated in MCPH patients have been identified so far (*MCPH1*-*20*), including genes encoding proteins associated with centriole biology or the mitotic spindle such as *ASPM* (MCPH5) and *WDR62* (MCPH2), the two most commonly mutated MCPH genes (Jayaraman, Kodani et al., 2016, Megraw, Sharkey et al., 2011, Saade, Blanco-Ameijeiras et al., 2018, Thornton & Woods, 2009). Centrioles are barrel-shaped, membrane-less organelles which behave as the major microtubule organizing centers (MTOCs) generating key cellular structures such as centrosomes, cilia and flagella. Structurally, centrioles consist of nine microtubule triplets organized around a cartwheel from which nine protein spokes emanate radially confering a 9-fold symmetric conformation that stabilizes the structure (Bettencourt-Dias & Glover, 2007). Centrioles typically duplicate once per cell cycle in a semiconservative fashion during the G1-to-S phase transition generating two fully functional centrosomes with two centrioles each. These structures recruit several proteins to polymerize a robust pericentriolar matrix (PCM) thereby allowing the generation of a bipolar spindle that ensures the correct segregation of chromosomes to both daughter cells thus avoiding chromosomal instability. Deregulation of centrosome dynamics typically results in variety of cell division defects ultimately affecting neural development (Saade et al., 2018).

Centriole duplication relies in the formation of a newborn pro-centriole in an orthogonal angle using the preexisting centriole as a biogenesis platform (Bettencourt-Dias & Glover, 2007). The core centriole duplication toolbox required for this process includes enzymatic components such as Polo-like kinase 4 (Plk4), which acts as a key regulator of centriole biogenesis (Bettencourt-Dias, Rodrigues-Martins et al., 2005, Habedanck, Stierhof et al., 2005), as well as structural components such as Sas4 (MCPH6, also known as Cpap or Cenpj)(Hung, Tang et al., 2000), Sas6 (MCPH14)(Leidel, Delattre et al., 2005) or Cep135 (MCPH8)(Ohta, Essner et al., 2002). Cep135 was originally identified as a component of the cartwheel with critical scaffold function in centriole structure and biogenesis (Ohta et al., 2002). Pioneer studies in *Drosophila* suggested minor defects in centriole duplication after *Cep135* knockdown (Carvalho-Santos, Machado et al., 2012, Dobbelaere, Josue et al., 2008, Mottier-Pavie & Megraw, 2009, Roque, Wainman et al., 2012), as well as mild perturbations of centriole structure in *Cep135* mutant spermatocytes (Roque et al., 2012). *Cep135* knockdown in chicken cell lines does not significantly alter centriole structure and organization, nor centriole biogenesis or cell proliferation, and only mild centrosome duplication defects arise (Inanc, Putz et al., 2013). In contrast, genetic knockdown by RNA interference of *CEP135* in human cell lines induced significant centriole duplication defects, altered spindle assembly and aberrant centriole structure (Lin, Chang et al., 2013). Interestingly, genetic analysis of a cohort of MCPH patients, identified a homozygous single base-pair deletion in exon 8 of human *CEP135* (also known as MCPH8) which resulted in a frameshift that produced a premature stop codon (Hussain, Baig et al., 2012). Patients bearing this mutation present severe microcephaly and cognitive deficits although the pathogenic mechanism leading to these defects are unclear.

In this work, we have generated a series of novel *Cep135*-deficient mouse models with different Cep135 protein levels by using CRISPR/Cas9-mediated direct editing of mouse embryos. Our data show that Cep135 inactivation in the mouse results in primary microcephaly accompanied by a p53-associated antiproliferative response in the developmental cortex. Genetic ablation of p53 prevents apoptosis during mid-gestation but does not prevent microcephaly caused by *Cep135* ablation in these mutant embryos. Lack of *Cep135* in fibroblasts expanded in vitro and neural progenitors in vivo results in several abnormalities in cell division in the presence of impaired centriole duplication. These defects are not seen in *Aspm* or *Cdk5rap2* (MCPH3)-mutant embryos suggesting that *CEP135* mutations likely cause primary microcephaly by specifically preventing proper dynamics of centrosome duplication in neural progenitors during development.

## Results

### Genetic depletion of *Cep135* promotes microcephaly in mice

We targeted the second exon of murine *Cep135* using two single guide RNAs (sgRNAs) generating a mutant sequence with a 278 bp deletion [*Cep135*(Δ278) allele lacking the initial ATG site] and a 8 bp deletion [*Cep135*(Δ8) allele] (Fig. 1a and Supplementary Fig. 1a). *Cep135*(Δ278/Δ278) mouse embryonic fibroblasts (MEFs) displayed reduced protein levels at the centrosome whereas no detectable Cep135 was present in *Cep135*(Δ8/Δ8) cells (Fig. 1b). Combination of these alleles in either homozygosity or heterozygosity allowed the generation of six cohorts of *Cep135*-mutant mice with different amounts of Cep135 protein, ranging from wild-type levels to intermediate levels in *Cep135*(Δ278/Δ278) homozygous mutants and *Cep135*(Δ278/Δ8) heterozygous mice, and no detectable protein in the *Cep135*(Δ8/Δ8) model (Fig. 1c).

**Figure 1.**
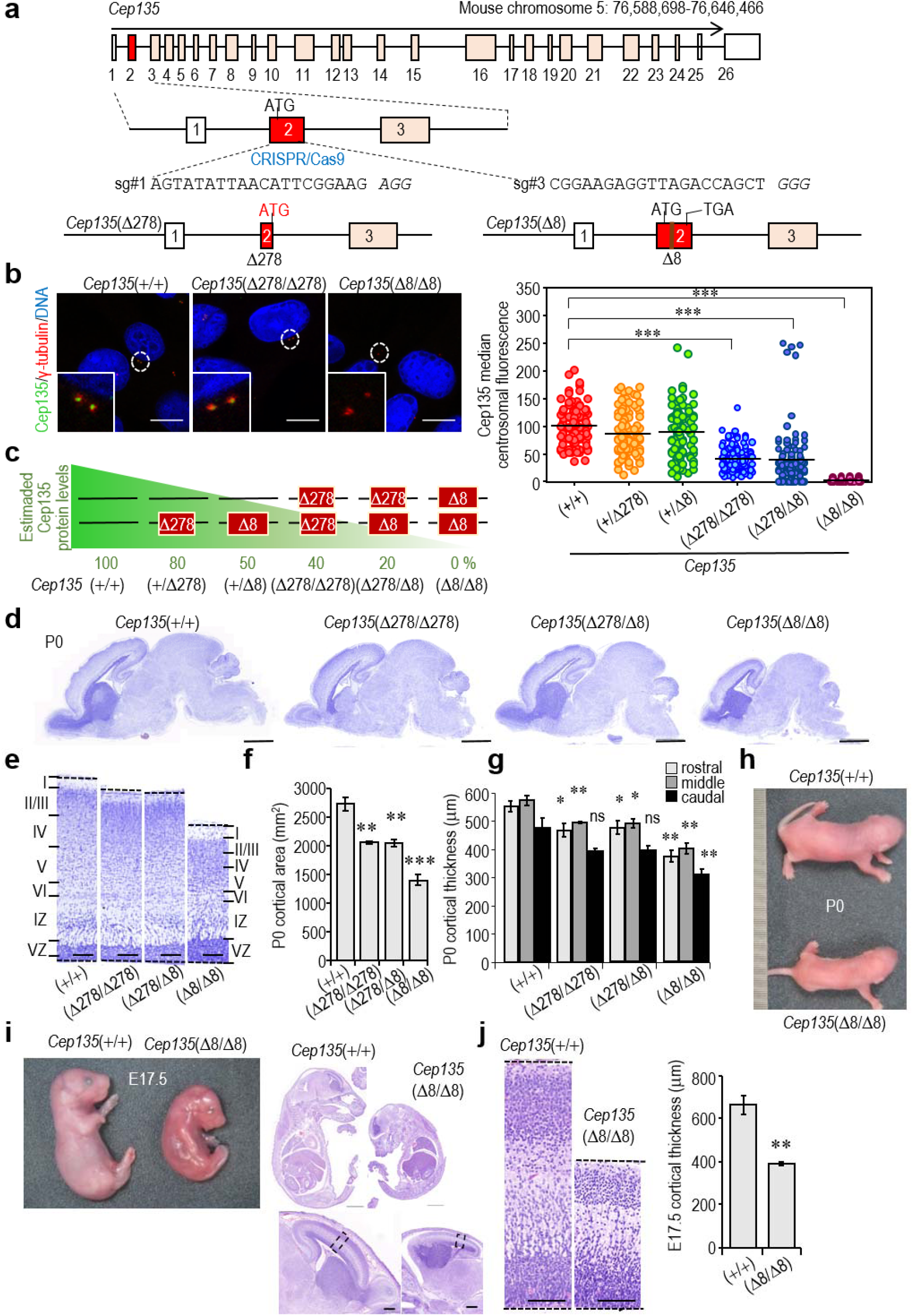
A novel model of microcephaly caused by *Cep135* ablation. **a**, Generation of the *Cep135*(Δ278) and *Cep135*(Δ8) by CRISPR/Cas9-mediated gene editing in mouse embryos. The *Cep135*(Δ278) allele lacks the initial ATG site and contains a second ATG that could generate a Cep135 truncated protein missing 89 aa of the N-terminal domain, a domain critical for microtubule binding. The Δ8 allele generates of a premature stop codon in exon 3. **b**, Confocal imaging of centrosomal loading of Cep135 (green) in control and mutant MEFs with different combinations of *Cep135*(Δ278) and *Cep135*(Δ8) mutant alleles. γ-tubulin is depicted in red, Cep135 in green, and DNA (blue) was stained with 4′,6-diamidino-2-phenylindole (DAPI). Scale bars: 10 μm. The plot to the right shows the quantification of Cep135 centrosomal protein levels (median fluorescence intensity in arbitrary units) in cells with the indicated genotypes. Horizontal bars depict the mean; ***, *P*<0.001 (Unpaired t-test with Welsh correction). **c**, A summary of the effect of the different *Cep135* alleles in the centrosomal loading of Cep135. **d**, Representative micrographs of sagital brain sections stained with Nissl at postnatal day 0 (P0) in neonate *Cep135* mutants. Scale bars: 1 mm. **e**, Histological Nissl staining of P0 cortices from mice with the indicated *Cep135* genotypes. Roman numerals indicate approximate cortical layers in P0 developing brains; IZ: intermediate zone; VZ ventricular zone. Scale bars: 100 μm. **f**, Quantification of cortical area in brain sections from *Cep135* mutant neonates (P0). **g**, Quantification of cortical thickness in rostral, medial, caudal aspects of P0 *Cep135* mutant mice versus *Cep135*(+/+) controls. **h**, Representative macroscopic images of *Cep135*(+/+) and *Cep135*(Δ8/Δ8) P0 neonatal pups. **i**, Representative macroscopic images of *Cep135*(+/+) and *Cep135*(Δ8/Δ8) E17.5 fetuses (left panel). The right panels show haematoxylin and eosin (H&E) staining of histological sagital sections from E17.5 *Cep135*-null and control fetuses. Scale bars: 1 mm (top) and 500 μm (bottom panel showing a higher magnification of the brain area). **j**, Representative images of H&E-stained sections of cortices from these E17.5 fetuses. Scale bars: 100 μm. The histogram to the right shows the quantification of cortical thickness of the medial aspect in these neocortices. Data in **f**,**g**,**j** represent mean ± SEM from 3 different mice or embryos; ns, non-significant; *, *P*<0.05; **, *P*<0.01; ***, *P*<0.001 (Student’s t-test).

These alleles allowed a comparison between *Cep135* gene dosage and the phenotype arising from different Cep135 protein levels in vivo. Both *Cep135*(Δ278/Δ278) and *Cep135*(Δ278/Δ8) newborns (postnatal day 0 or P0) displayed mild microcephaly (∼24.5% and 24.7% reduction in cortical area, respectively; Fig. 1d-f), whereas homozygous *Cep135*(Δ8/Δ8) P0 mice showed severe microcephaly as showed by a reduced neocortical area (48.5% cortical area reduction; Fig. 1f) and reduced neocortical thickness in the rostral, medial and caudal aspects of the neonatal neocortices (Fig. 1g). Full depletion of Cep135 in the *Cep135*(Δ8/Δ8) model resulted in smaller full body size (Fig. 1h) and perinatal lethality with all mutant mice dying within a few hours after birth. Histological examination of *Cep135*-null intraperitoneal organs revealed simplified and underdeveloped intestinal and stomachal mucosae with no evidences of recent lactation, immature or underdeveloped lungs or retinal abnormalities (Supplementary Fig. 1b,c). Detailed analysis of the lungs revealed small, immature alveolae with lower levels of Surfactant Protein-C (Supplementary Fig. 1d), suggestive of respiratory distress. *Cep135*(+/Δ278) and *Cep135*(+/Δ8) heterozygous mice exhibited no significant differences in the centrosomal levels of Cep135 (Fig. 1b) and, accordingly, no microcephaly, body size reduction or other phenotypic defects were observed (data not shown).

Examination of *Cep135*(Δ8/Δ8) fetuses at embryonic day 17.5 (E17.5) confirmed a smaller overall fetal body size and severe microcephaly when compared to their wild-type littermates (Fig. 1i), with reduced thickness of the developing neocortex (Fig. 1j). The reduction in head and brain weight was more pronounced than the reduction in body weight (Supplementary Fig. 1e) suggesting that *Cep135* depletion may promote microcephaly in a gene dose-dependent manner in mice.

### p53 mediates cell death of Cep135-deficient cortical neural progenitors

To gain insight into the molecular pathways deregulated during cortical development in *Cep135*-mutant embryos, we subjected developing cortices from E11.5 and E14.5 *Cep135*(Δ8/Δ8) and control *Cep135*(+/+) embryos to RNAseq analysis. In addition, E14.5 cortices were disaggregated and cultured to form neurospheres and RNA from these in vitro cultures was analyzed after 12h and 24h (Figure 2a). Combined analysis of transcriptomic profiles from these samples segregated three main groups corresponding to E11.5 cortices, E14.5 cortices and cultured cells (12h and 24 h time points; Figure 2b). Differential expression between *Cep135*-null and control samples in each group (Supplementary Figure 2a) suggested a defect in neural differentiation and function in *Cep135*-deficient samples both in vivo at E11.5 (Figure 2c, Supplementary Figure 2b and Supplementary Table 1) and E14.5 (Supplementary Figure 2c and Supplementary Table 2), as well as in the neurosphere formation assay (Figure 2d, Supplementary Figure 2c and Supplementary Tables 3 and 4). Interestingly, the p53 pathway was deregulated in these samples with significant upregulation of multiple p53 target genes, and a significant enrichment of deregulated transcripts with p53 binding sites in their promoters (Figure 2c,d, Supplementary Figure 2c and Supplementary Tables 5-8). The p53 target and cell cycle inhibitor *Cdkn1a* (p21^Cip1^) was among the most upregulated genes in these samples (Supplementary Table 9).

**Figure 2.**
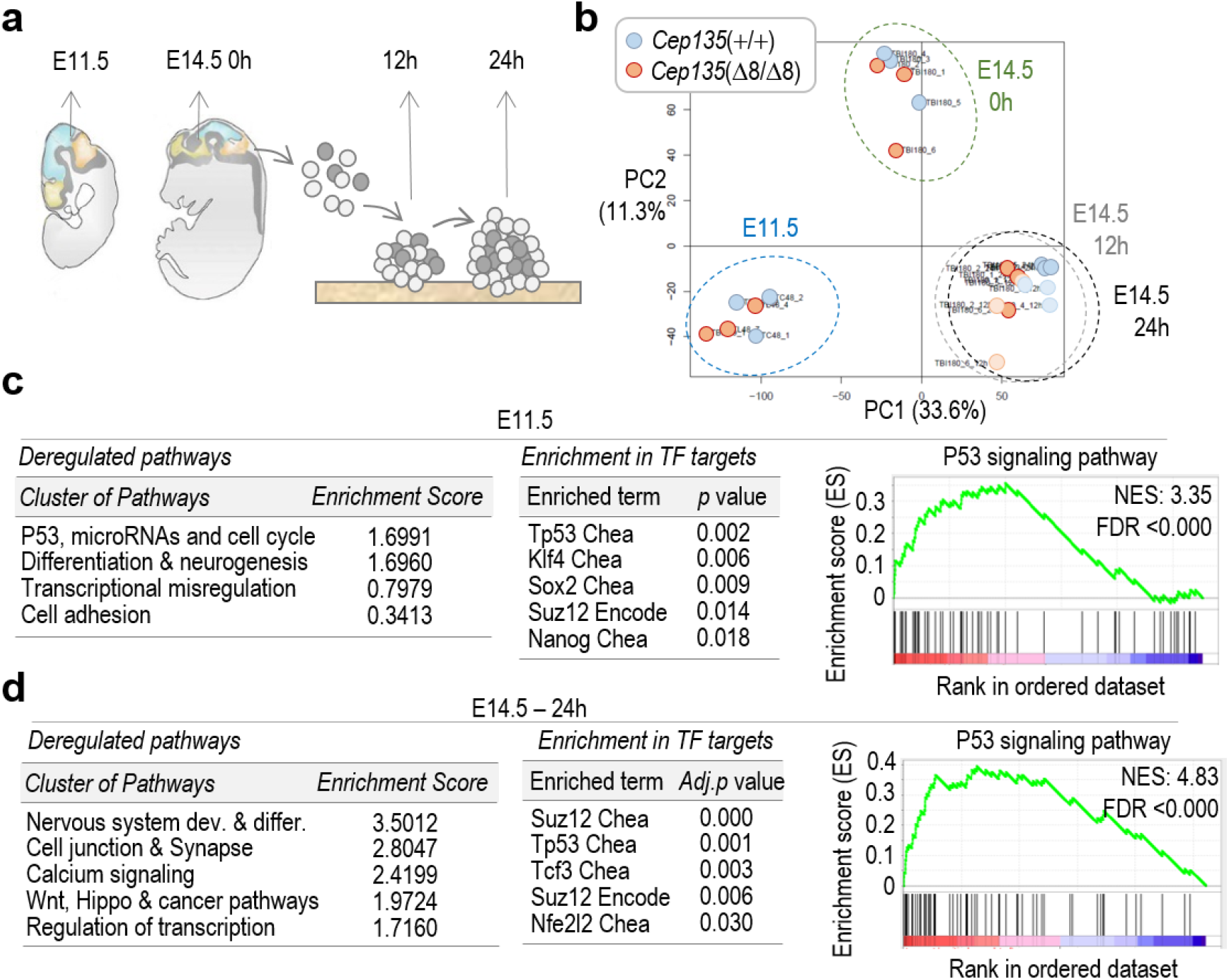
p53-mediated response to lack of Cep135 in developing brains and cultured neurospheres. **a**, Schematic representation of the *Cep135*(+/+) and *Cep135*(Δ8/Δ8) samples selected for RNA sequencing analysis, including E11.5 and E14.5 brains, as well as neurospheres from E14.5 brains cultured for 12 and 24 h. **b**, Principal component analysis of the transcriptomic profiles from the indicated samples. Values in PC labels show the percentage of explained variance for PC1 and PC2. **c**, Major pathways deregulated and enrichment in transcription factor (TF) targets in E11.5 samples. The enrichment in transcripts involved in the p53 pathway is shown to the right. **d**, Major pathways deregulated and enrichment in transcription factor (TF) targets in E14.5 samples cultured during 24 h to form neurospheres. The enrichment in transcripts involved in the p53 pathway is shown to the right. See Supplementary Tables 1-9 for additional details.

At the protein level, p53 was highly induced in several tissues in E11.5 *Cep135*(Δ8/Δ8) embryos (Fig. 3a). This response was accompanied by a significant number of apoptotic cells (as detected by active caspase 3) in the developing brain and the rest of the body, especially dorsal telencephalon cortical cells and hematopoietic progenitors in the embryonic liver. At E14.5, *Cep135*(Δ8/Δ8) embryos were smaller and apoptotic cells were observed in the developing neocortex but were almost absent in the rest of the embryonic tissues (Fig. 3b). Immunostaining for cleaved caspase 3 in whole embryos suggested that apoptosis was especially evident in the developing neocortex at this stage (Fig. 3c). Additional immunofluorescence studies for Sox2 and Tuj1 to detect apical progenitors and neuroblasts, respectively, showed apoptotic cell death in both progenitor and neuroblast lineages in the E14.5 developing neocortex (Fig. 3d). E11.5 and E14.5 *Cep135*(Δ8/Δ8) embryos also displayed reduced number of proliferating (as defined by phospho-histone H3-positive mitoses) cells in several tissues (Supplementary Fig. 3a,b), suggesting that both early apoptosis and defective proliferation may contribute to reduced body size in these mutant Cep135-deficient embryos. No evidence of apoptotic cells was observed in *Cep135*-mutant embryos at E17.5 (Supplementary Fig. 3c), suggesting clearance of apoptotic cells and adaptation to the lack of Cep135 during the later stages of development.

**Figure 3.**
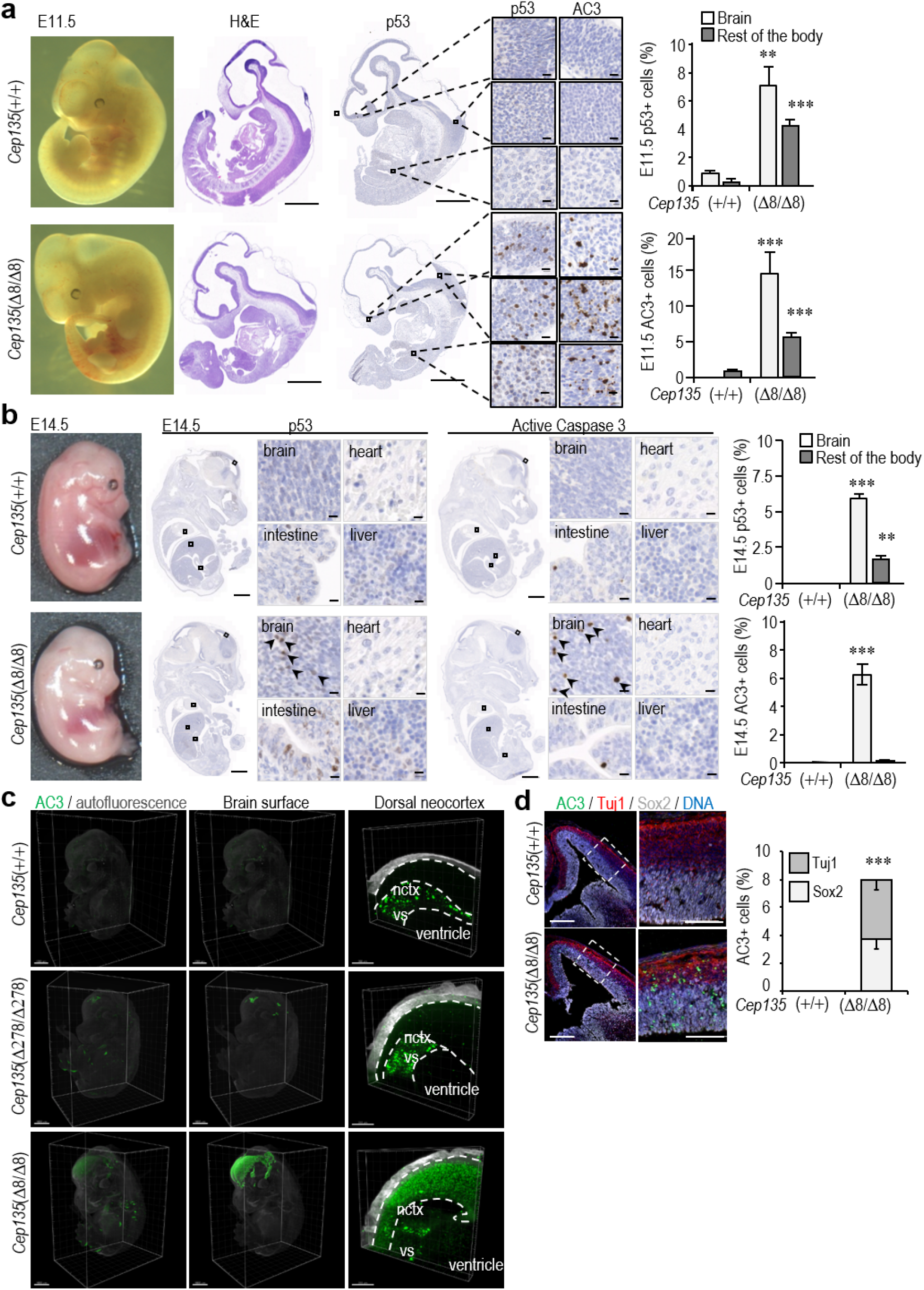
Cep135 deficiency induces p53-dependent apoptosis in neural progenitors. **a**, Bright-field macroscopic (left) and haematoxylin and eosin staining (H&E, middle) images of E11.5 *Cep135*(+/+) and *Cep135*(Δ8/Δ8) mouse embryos. Immunohistochemical staining for p53 and active caspase 3 (AC3) in the same samples including insets at higher magnification (p53 left, AC3 right) from the specific areas [forebrain (up), medullary hindbrain (middle) and liver (bottom)]. The quantification of p53 and AC3-positive cells in brain and rest of the body of *Cep135*-mutant and control embryos is shown in the right histograms. Scale bars: 1 mm (whole embryo sections), 10 μm (insets). **b**, Brightfield macroscopic images (left), immunohistochemical staining (middle) and quantification (right) of p53 and AC3-positive cells in brain and rest of the body of E14.5 *Cep135*-mutant and control embryos. Scale bars: 1 mm (whole embryo sections), 100 μm (lower microscopic insets). Representative positive cells are indicated by arrows. **c**, Wholemount 3D immunofluorescence of E14.5 *Cep135*(+/+), *Cep135*(Δ278/Δ278) and *Cep135*(Δ8/Δ8) mouse embryos stained for AC3 (green). Gray color depicts tissue autofluorescence in the 488 channel. Scale bars: 1 mm (whole embryos); 200 μm (lower microscopic insets). Nctx: neocortex; vs: ventricular surface. Note that green, unspecific positive signal in the vs of *Cep135*(+/+) samples corresponds to secondary antibody aggregates. **d**, Immunostaining for AC3, Sox2 and Tuj1 in 14.5 neocortex from *Cep135*-null and control mice. Scale bars: 250 μm (left), 100 μm (right). The bottom histogram shows the quantification of AC3-positive cells in these samples. In **a, b, d** data are mean SEM from 3 different embryos; **, *P*<0.01; ***, *P*<0.001 (Student’s t-test).

We next tested the functional relevance of p53 in the phenotype of *Cep135*-deficient embryos by interbreeding *Cep135*(+/Δ8) with p53-deficient mice. Lack of p53 protein (Fig. 4a) prevented apoptosis in *Cep135*(Δ8/Δ8); *Tp53*(–/–) E11.5 (Supplementary Fig. 3d) and E14.5 (Fig. 4b) mouse embryos. Intriguingly, p53 loss did not rescue the reduced thickness of the neocortex of *Cep135*-null embryos (Fig. 4b). In fact, whereas we did not observe increased lethality of E11.5 *Cep135*(Δ8/Δ8); *Trp53*(–/–) double mutant embryos, we only recovered one double mutant embryo alive out of 39 E14.5 embryos derived from 4 matings between *Cep135*(+/Δ8); *Trp53*(–/–) males and *Cep135*(+/Δ8); *Trp53*(+/–) females; with two additional double mutant dead embryos being reabsorbed (Supplementary Fig. 3e), suggesting premature embryonic death and embryonic reabsorption at mid-gestation. This E14.5 *Cep135*(Δ8/Δ8); *Tp53*(–/–) double mutant embryo presented profuse microcephaly with severe cortical malformations such as abundant aberrant mitotic figures, perturbed laminar layering of neural progenitors and abnormal neocortical architecture organization (Fig. 4c). Additionally, *Cep135*(Δ8/Δ8); *Tp53*(+/–) double mutant embryos also presented clear cortical malformations, together with severe cortical dysplasia and heterotopias within the E14.5 neocortex. Together, these data suggest that Cep135 loss promotes whole-body proliferative defects during mouse embryonic development, and a p53-dependent cell death of neural progenitors and neuroblasts resulting in severe microcephaly.

**Figure 4.**
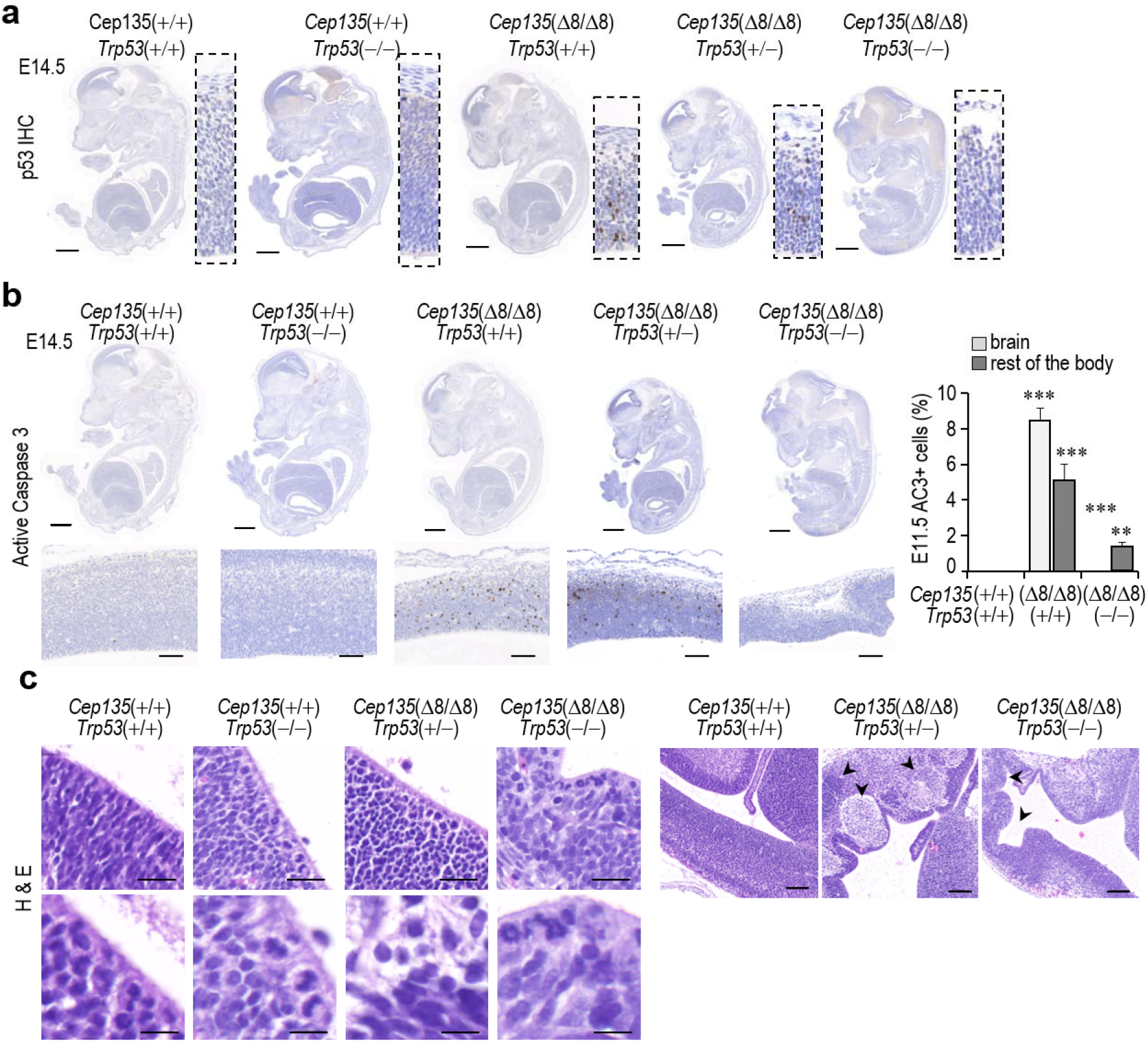
Genetic ablation of *Trp53* rescues cell death but not microcephaly in *Cep135* knockout embryos. **a**, Immunohistochemical staining for p53 (brown, whole embryos and insets) in *Cep135*(+/+) and *Cep135*(Δ8/Δ8) E14.5 embryos in a *Trp53*(+/+), *Trp53*(+/–) and *Trp53*(–/–) genetic background. Scale bars: 1 mm. **b**, Immunohistochemical staining for active caspase 3 (AC3, brown) of E14.5 embryos with the indicated genotypes from crosses between *Cep135* and *Trp53*-mutant mice. Higher magnification insets below depict a representative area of the medial section of the neocortex. Scale bars: 1 mm (whole embryo sections), 100 μm (insets). The percentage of AC3-positive cells in brains and bodies from E11.5 *Cep135* and *Trp53*-double mutant embryos is shown to the right. Data are mean SEM from 3 different embryos; **, *P*<0.01; ***, P<0.001 (Student’s t-test). **c**, H&E staining of the medial aspect of the neocortex of E14.5 embryos with the indicated *Cep135* and *Trp53* genotypes. Scale bars: 20 μm (upper left panels), 10 μm (lower panels). 100 μm (panels to the right). Arrowheads indicate cortical malformations and heterotopias.

### Lack of Cep135 induces multiple defects in centrosome dynamics and chromosomal instability

To gain a deeper insight about the cellular alterations generated by Cep135 loss, we initially screened for centrosome or cytoskeleton defects in Cep135 deficient MEFs. Whereas two foci of γ-tubulin [a major component of the pericentriolar material (PCM)] were detected in most control cells positive for cyclin A (a marker of late S-G2), only one γ-tubulin structure was found in the majority of cyclin A^+^ cells derived from *Cep135* knockout embryos (Fig. 5a). For a more detailed analysis of cell cycle stages, we made use of proliferating cell nuclear antigen (PCNA), a DNA replication processivity factor whose nuclear pattern changes depending of the period of the S-phase, and phospho-histone H3 (PH3), a modification that correlates with chromosome condensation during late G2 and mitosis. Whereas both control and *Cep135*-mutant cells displayed a single centrosome in G1, co-immunostaining of γ-tubulin with PCNA revealed efficient centrosome duplication in control *Cep135*(+/+) but not in mutant *Cep135*(Δ8/Δ8) cells (Fig. 5b). In about 50% of *Cep135-*null cells, a single γ-tubulin spot was present from G1 to late G2, inducing monopolar spindles during mitosis (Fig. 5b). Interestingly, the remaining fraction of *Cep135* knockout cells that contained two centrosomes presented additional defects, such as centrosomal asymmetry in the maturation of centrosomes when stained with Centrin (a centriolar protein) and the PCM component γ-tubulin. This defect typically affected the daughter, newly generated centrosome, and not the mother centrosome which is characterized by the presence of Outer Dense Fiber Protein 2 (Odf2; Fig. 5c).

**Figure 5.**
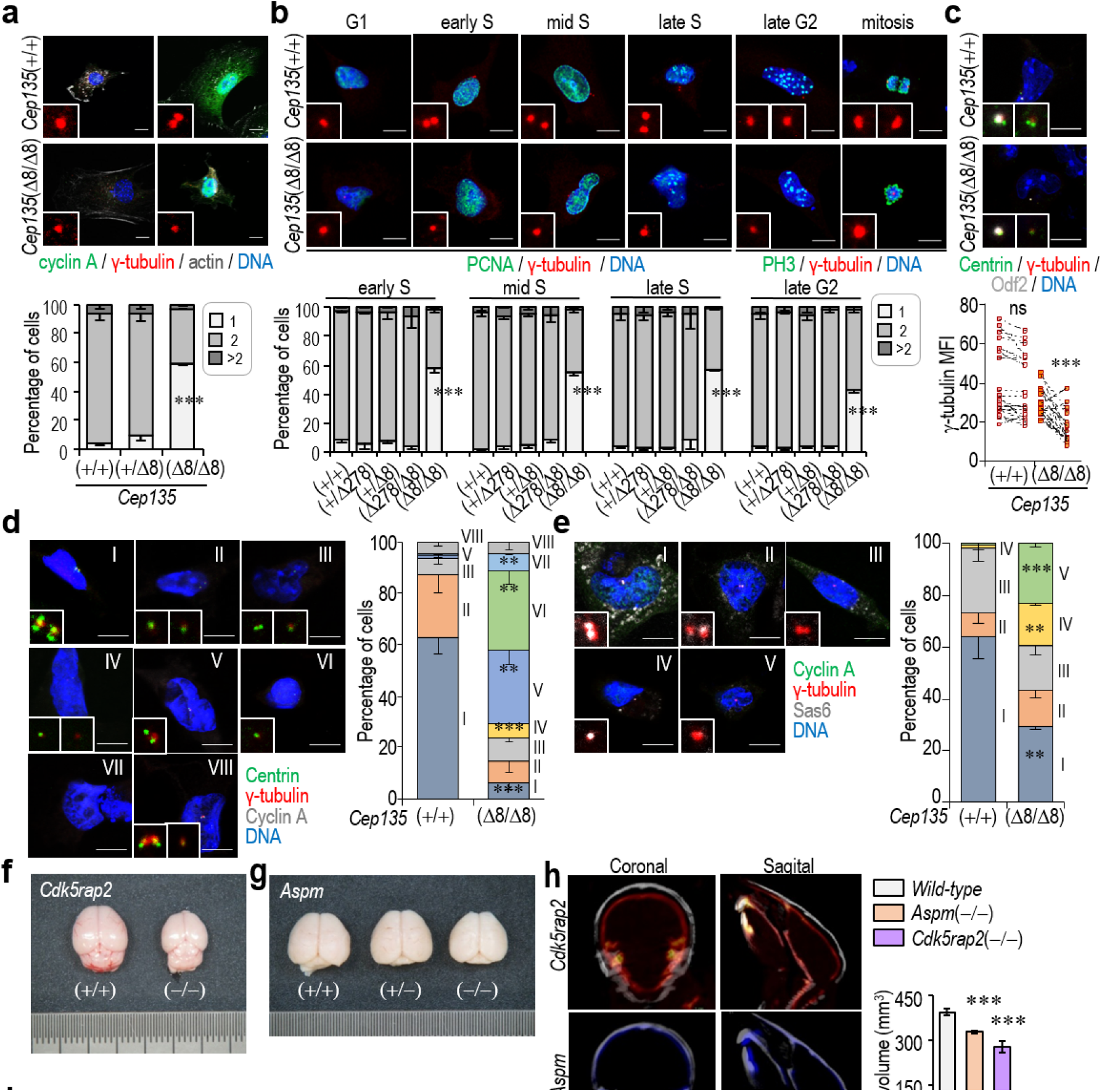
*Cep135*-deficient fibroblasts display centrosome dynamics defects, aberrant mitoses and chromosomal instability. **a**, Confocal imaging of E14.5 *Cep135*-mutant and control MEFs stained with phalloidin (actin cytoskeleton, grey), DAPI (DNA, blue), antibodies against cyclin A (green) and γ-tubulin (red). The bottom histogram shows the percentage of cells with 1, 2 o more than 2 γ-tubulin spots. **b**, Immunostaining of *Cep135*-mutant and control MEFs with antibodies against PCNA (green, left group) or phospho-histone H3 Ser10 (PH3; green, right group), antibodies against γ-tubulin (red) or DAPI (DNA, blue). The bottom histograms show the percentage of cyclin A+ cells with 1, 2 o more than 2 γ-tubulin spots in the different phases of the cell cycle, as determined by PCNA and PH3 staining. **c**, Immunostaining with antibodies against centrin (green), γ-tubulin (red), and Odf2 (Cenexin, a marker of the mother centrosome; grey). The bottom panel shows the quantification of γ-tubulin median centrosomal fluorescence intensity (MFI). Slashed lines link two centrosomes from the same cell; higher slope of the line indicates higher centrosomal asymmetry of γ-tubulin fluorescence intensity between both centrosomes. **d**, Confocal imaging of E14.5 *Cep135*(Δ8/Δ8) MEFs stained with DAPI (DNA, blue) and antibodies against centrin (green), γ-tubulin (red), cyclin A (grey). All cells depicted are cyclin A+. Group I: Two γ-tubulin spots; 2 centrin doublets; II: Two γ-tubulin spots; 2 centrin singlets; III: Two γ-tubulin spots; 1 centrin doublet + 1 centrin singlet; IV: Two γ-tubulin spots; 1 centrin doublet + 1 centrin singlet; V: 1 γ-tubulin spot; 1 centrin doublet; VI: 1 γ-tubulin spot; 1 centrin singlet; VII: acentrosomal; VIII: >2 γ-tubulin spots. The histogram to the right shows the percentage of cells in each group. **e**, Confocal imaging of E14.5 *Cep135*(Δ8/Δ8) MEFs stained with DAPI (DNA, blue) and the indicated antibodies. All cells depicted are cyclin A+. Group I: 2 γ-tubulin spots; 2 Sas6 singlets; II: 2 γ-tubulin spots; 1 Sas6 singlet + no Sas6; III: 2 γ-tubulin spots; no Sas6 – no Sas6; IV: 1 γ-tubulin spot; 1 Sas6 singlet; V: 1 γ-tubulin spot; no Sas6. The histogram to the right shows the percentage of cells in each group. **f**, Macroscopic images of *Cdk5rap2*(+/+) and *Cdk5rap2*(–/–) P30 brains. **g**, Macroscopic images of *Aspm*(+/+) and *Aspm*(–/–) P30 brains. **h**, Micro-CT imaging of wild-type (white), *Aspm-null* (blue) and *Cdk5rap2-null* (red) P30 skulls in coronal and sagital views. The quantification is shown in the histogram to the right. **i**, Confocal imaging of E14.5 *Aspm*-mutant, *Cdk5rap2*-mutant and control MEFs stained with DAPI (DNA, blue) and antibodies against cyclin A (gray), γ-tubulin (red) and centrin (green). The side histogram shows the percentage of cyclin A+ cells with 1 or 2 γ-tubulin spots. In **a**,**b**,**c**,**d**,**e**,**i** scale bars: 10 μm. In **a-e**,**h**,**i** data are mean ± SEM from 3 different primary cultures from 3 different E14.5 embryos; n>150 cells/each independent condition; **, *P*<0.01; ***, *P*<0.001 (Student’s t-test).

Combined immunofluorescence with centrin and γ-tubulin in cyclin A-positive cells indicated that centrosome duplication was compromised in *Cep135*(Δ8/Δ8) cells (Fig. 5d, group V). In addition, these mutant cultures were enriched in cells with only one centrosome and one centriole, suggestive of both centrosome duplication and centriole assembly defects (Fig. 5d, VI), as well as in cells lacking centrosomes (acentrosomal; Fig.5, VII). Similar results were obtained in late G2 PH3+ cells, confirming that centriole defects caused by Cep135 depletion were also sustained during the rest of the cell cycle (Supplementary Fig. 4a). Immunostaining of S/G2 cells (Cyclin A^+^) for PCM (γ-tubulin) and cartwheel proteins (Sas6) revealed a significant increase in cells with only one centrosome devoid of cartwheel (Fig. 5e; group V), suggesting additional defects in cartwheel assembly.

Despite the frequency of mutations in centriolar *MCPH* genes, the lack of comparative studies makes it unclear to what extent these mutations result in similar cellular defects. We therefore decided to generate new strains of mice mutant in *Cdk5Rap2* (*MCPH3*) or Aspm (*MCPH5*, mutated in 60% of MCPH patients) using CRISPR/Cas9-mediated gene editing in mouse embryos. Both *Aspm*(–/–) and *Cdk5rap2*(–/–) mutant mice displayed microcephaly by P30 (Fig. 5f-h). Interestingly, MEFs derived from *Aspm* and *Cdk5rap2*-mutant embryos presented normal centrosome duplication during S phase (Fig. 5i) as opposed to *Cep135*-null MEFs, suggesting that centrosome duplication defects are a specific feature generated by the absence of Cep135.

The presence of two properly matured and separated centrosomes is critical for the establishment of a bipolar spindle and correct chromosome segregation. Time-lapse microscopy analysis of *Cep135*(Δ278/Δ278) or *Cep135*(Δ278/Δ8) cultures showed no alteration in the duration of mitosis. However, homozygous cells for the *Cep135*(Δ8) mutation displayed significantly increased duration of mitosis, in agreement with abundant monopolar spindles (Supplementary Fig. 4b,c). These defects were accompanied by increased DNA content (4n and over-4n) and aneuploidy in a *Cep135* gene dose dependent manner (Supplementary Fig. 4d,e). All together, these results suggest that loss of Cep135 leads to defective centriole dynamics, ultimately inducing chromosomal instability, at least in fibroblasts.

### Perturbed centrosomal and mitotic dynamics in *Cep135*-deficient neural progenitors

We next examined the effect of lack of Cep135 in cortical neural progenitors isolated from E14.5 neocortices and cultured to generate primary neurospheres. In agreement with the observations in fibroblasts, *Cep135*-deficient neurospheres displayed centrosome duplication defects (Fig. 6a), accumulation of p53 (Fig. 6b) and apoptotic cell death (Fig. 6c). *Cep135*-deficient primary neural progenitors also displayed a significant defect in their self-renewal capacity as scored by the limiting dilution assays (Fig. 6d).

**Figure 6.**
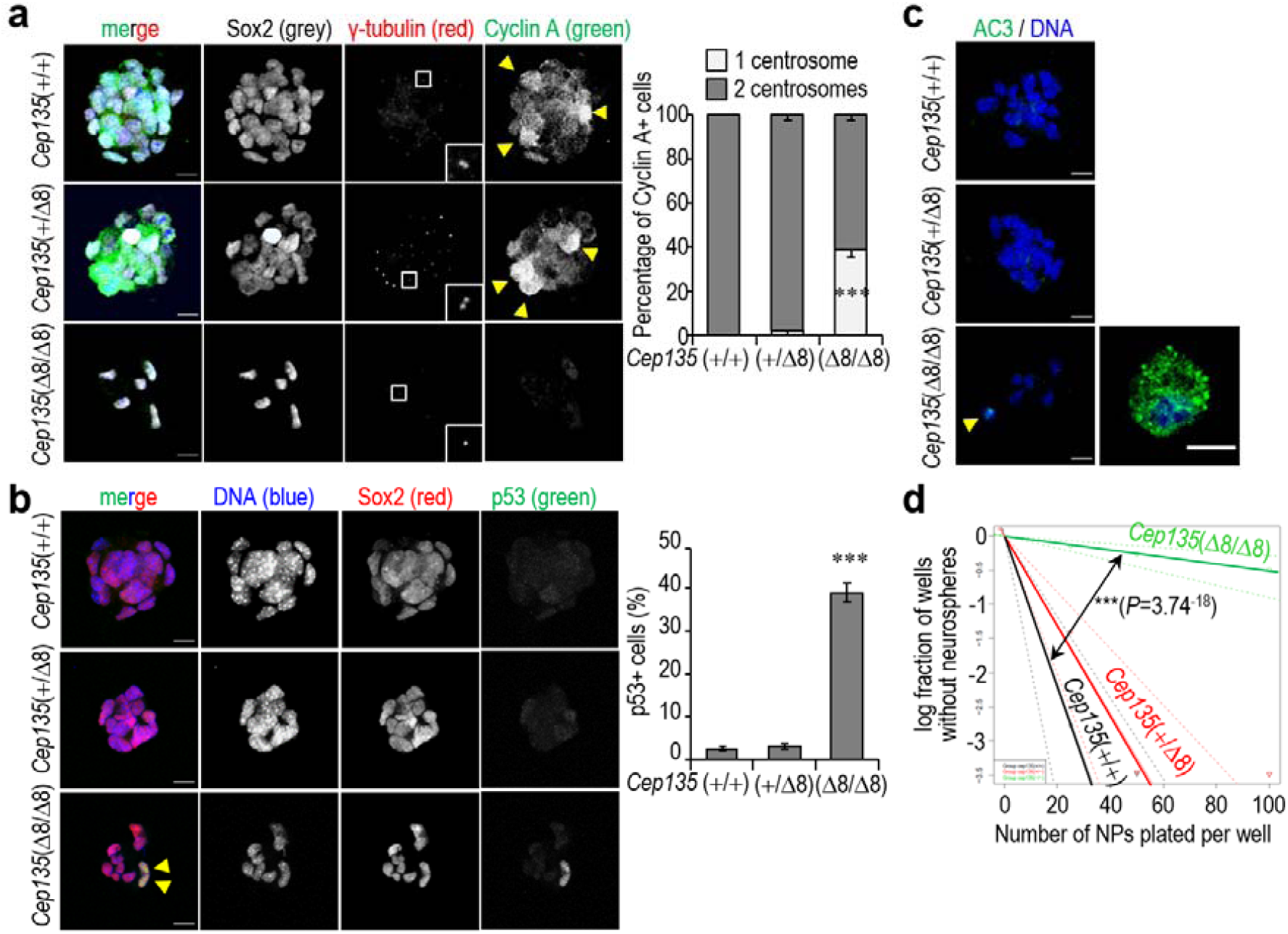
Neurosphere formation and self-renewal abilities in *Cep135*-mutant neural progenitors. **a**, Confocal imaging of incipient neurospheres derived from E14.5 embryonic cortices after staining with the indicated antibodies. The percentage of cells positive for cyclin A with one or two centrosomes is shown to the right. **b**, Immunostaining of primary neurospheres from E14.5 embryos with Sox2 and p53 antibodies. The percentage of p53+ cells is shown to the right. **c**, Confocal imaging of primary neurospheres stained with antibodies against active caspase 3 (AC3, green). DAPI (DNA) is shown in blue. **d**, Self-renewal ability of neural progenitors with the indicated genotypes as determined by limiting dilution assays. In **a**,**b**,**c**, scale bars are 10 μm, and arrowheads indicate Cyclin A+ cells (**a**), p53+ cells (**b**) or apoptotic cells (**c**). In **a**,**b** data are mean ± SEM. ***, *P*<0.001 (Student’s t-test in **a**,**b**, and Chi-squared in **d**).

Unfortunately, these defects did not allow to establish cultures of *Cep135*(Δ8/Δ8) neural progenitors for further assays of centrosome dynamics in vitro. We therefore looked for centrosome alterations in situ. The reduced cortical thickness observed at birth (Fig. 1) was also present in these mutant mice at E14.5 and specifically affected the Sox2-positive apical progenitor (AP) and the Tbr2-positive intermediate progenitor (IP) cell populations within the developing neocortex (Fig. 7a). No significant differences were observed in the orientation of the mitotic spindle in dividing cells of the AP layer in *Cep135*-deficient embryos (Supplementary Fig. 5a). However, while most mitoses were bipolar in *Cep135*(+/+) embryos, *Cep135*(Δ8/Δ8) embryonic sections revealed frequent mitotic defects including monopolar or acentrosomal mitotic spindles in addition to apoptotic figures (Fig. 7b and Supplementary Fig. 5b). Maturation of centrosomes was impaired as detected by overall reduced γ-tubulin staining in APs of Cep135-deficient embryos (Fig. 7c,d). *Cep135*(Δ8/Δ8) APs also presented asymmetric centrosomes with one of the two centrosomes in the poles with dim or almost absent γ-tubulin signal and asymmetric immunostaining of the distal appendage mother centriole protein Odf2 (Fig. 7c). Electron microscopy (TEM) analysis of centrioles in APs showed an aberrant axial structure in *Cep135*(Δ8/Δ8) embryos, with lack or abnormal number of microtubule triplets, reduced centriolar diameter in some centriolar structures and lack of a clear nine-fold symmetric conformation (Fig. 7e). All together, these results suggest that alteration of Cep135 protein levels lead to defective centriolar duplication and dynamics in neural progenitors, accompanied by p53-dependent cell death and microcephaly.

**Figure 7.**
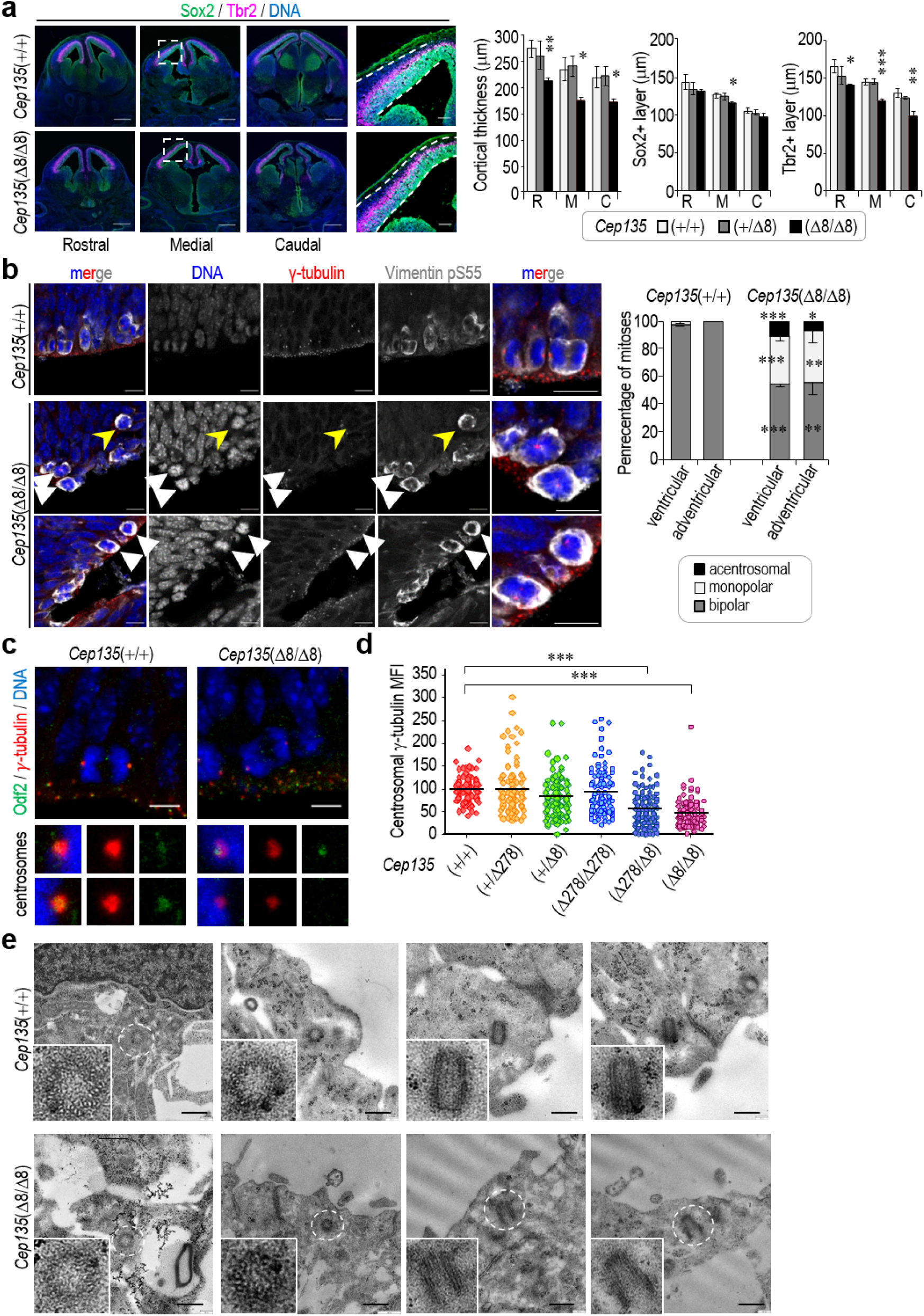
Aberrant centriole structure and organization in *Cep135*-deficient neural progenitors. **a**, Confocal imaging of cryosections of the rostral, medial and caudal aspects of E14.5 *Cep135*-mutant and control embryonic brains immunostained with antibodies against Sox 2 (green), Tbr2 (red) and with DAPI (DNA, blue); Scale bar: 1 mm (left panels) and 100 μm (insets). The histograms to the right show the quantification of the thickness (left), Sox-positive cells (middle) and Tbr2-positive cells (right) in the neocortices with the indicated genotypes in rostral (R), medial (M) and caudal (C) regions. **b**, Immunofluorescence with the indicated antibodies in developing E14.5 neocortex from embryos with the indicated genotypes. Control samples show typical bipolar spindles, whereas monopolar spindles (white arroheads in middle panels), acentrosomal spindles (yellow arrowheads) and asymmetric centrosomes (arrowheads in bottom panels) are observed in *Cep135*-mutant samples. Scale bars, 25 μm. The histogram on the right shows the quantification of polarity in mitotic spindles (monopolar, bipolar or acentrosomal) in ventricular or abventricular mitoses of E14.5 Cep135-mutant and control neocortices. In **a**,**b** data are mean ± SEM. *, *P*<0.05; **, *P*<0.01; ***, *P*<0.001 (Student’s t-test). **c**, Immunostaining with the indicated antibodies of anaphases in the ventricular surface of E14.5 *Cep135*-mutant and control embryos. Higher magnification images of mitotic centrosomes are displayed in the bottom panels. Scale bar: 10 μm. **d**, Quantification of median fluorescence intensity (MFI; arbitrary units) of γ-tubulin at the centrosome from NPs of the indicated genotypes. ***, *P*<0.001 (Unpaired t-test with Welsh correction). **e**, Transmission electron micrographs of E14.5 *Cep135*-mutant and control embryonic neocortices, showing representative pictures of centrioles in longitudinal and transversal view contained in the first layer of APs of the ventricular surface. Scale bars: 200 nm.

## Discussion

Despite the identification of at least 20 *MCPH* genes causative of primary microcephaly, the cellular basis underlying the neurodevelopmental abnormalities present in patients with most of these specific mutations is still elusive. Interestingly, more than half of *MCPH* gene products are known to be located at the centrosome or spindle poles, pointing to centrosome dysfunction as one of the main causes of MCPH (Jayaraman, Bae et al., 2018, Saade et al., 2018). The altered mitotic behavior commonly observed in MCPH usually comprises excessive asymmetric cell division of neural progenitors during brain development, impaired neural differentiation or cell death of progenitors, which accounts for a reduced pool of progenitors during development, reduced neuronal output and, as a result, microcephaly (Barbelanne & Tsang, 2014, Jayaraman et al., 2018).

Biallelic single base-pair deletions of the human *CEP135* gene have been reported in MCPH patients with significant microcephaly concomitant with severe retardation and communication deficits (Farooq, Fatima et al., 2016, Hussain et al., 2012). Previous reports included Cep135 in a group of centriolar proteins, including Sas4 and Sas6, thought to be essential for centriole assembly (Hung et al., 2000, Jayaraman et al., 2018, Leidel et al., 2005). Several studies have delineated a Wdr2-Aspm-Sas4-Sas6 pathway in which these MCPH-associated proteins recruit each other sequentially to the centrosome, thereby enabling centriole duplication to occur (Carvalho-Santos, Machado et al., 2010, Gonczy, 2012, Jayaraman et al., 2018). Cep135 interacts with Sas6 as well as tubulin participating in the interaction between the cartwheel and microtubule triplets (Hilbert, Noga et al., 2016).

Germline ablation of *Sas4* in the mouse results in severe defects in centriole biogenesis and duplications, leading to embryonic death as soon as E8.5 (Bazzi & Anderson, 2014). Conditional genetic depletion of these core structural proteins in later stages of embryonic development alter brain neurodevelopment by promoting apical neural progenitors (AP) detachment from the ventricular surface which results in spindle orientation randomization, ectopic proliferation of APs and progressive loss of centrioles concomitant with p53 activation and apoptotic death (Insolera, Bazzi et al., 2014). Sas4-null cells lack centrosomes and primary cilia; however, acentriolar spindle poles are assembled and these mutant cells do not display obvious defects in chromosome segregation (apart from slightly prolonged mitosis) or cell cycle profiles (Bazzi & Anderson, 2014). Similarly, genetic ablation of *SAS6* in human cells results in lack of centrioles (McKinley & Cheeseman, 2017). Whether similar requirements apply to mammalian Cep135 however is less clear. Pioneering *in vitro* studies of *Cep135* knockout using homologous recombination in vertebrate cell lines reported no significant alterations of centriole structure, dynamics or cell division, although a small decrease in centriole numbers and increase in the frequency of monopolar spindles was observed (Inanc et al., 2013). In contrast to these reports, we observed that complete genetic depletion of *Cep135* in vivo induces significant defects in centriole duplication, leading to frequent apoptotic cell death in several tissues by mid gestation. These defects are more pronounced and lasted longer in neural progenitors at later embryonic stages ultimately leading to microcephaly. As opposed to *Sas4*-knockout mice, which die before mid-gestation (Bazzi & Anderson, 2014, Insolera et al., 2014), *Cep135*-deficient mice progressed through embryonic development until birth, hence indicating that unlike Sas-4, Cep135 is dispensable for early embryonic development. The abnormal cell division of neural progenitors at mid gestation in the absence of Cep135 was companied by a p53 transcriptional response and p53-dependent apoptotic cell death. Ablation of the *Trp53* gene prevented apoptotic cell death but did not rescue the developmental defects caused by *Cep135* ablation. This is in contrast to previous microcephaly models in which the developmental delay and microcephaly caused by mutations in Sas4 (Bazzi & Anderson, 2014, Insolera et al., 2014), Kif20 (Little & Dwyer, 2019) or Cep63 (Marjanovic, Sanchez-Huertas et al., 2015) were rescued after *Trp53* ablation. The reasons for these differences are unclear but suggest that *Cep135* ablation may alter additional p53-independent pathways in the developing cortex.

Taken together, these data indicate that Cep135 depletion promotes centrosome duplication defects and mitotic aberrations accompanied with p53 activation and defective production of neurons during brain development. The resulting microcephaly (25% decrease of cortical area in Cep135-mutant newborns) can be considered as severe taking into consideration the ratio between human and murine encephalization, similar to other genetic mouse models based on *MCPH* gene mutations. The mechanism underlying *Cep135*-null phenotypes, i.e. defective centriole duplication, may however be relatively specific. Previous studies have shown that *Aspm* (which accounts for approximately 60% of human cases of MCPH) depletion in mice promotes mild microcephaly and germline defects and affects spindle orientation and astral microtubule dynamics (Fish, Kosodo et al., 2006, Gai, Bianchi et al., 2016, Pulvers, Bryk et al., 2010). Similarly, *Cdk5rap2* loss generates severe microcephaly and dwarfism by affecting cell division causing cell death of neural progenitors during development (Lizarraga, Margossian et al., 2010). Interestingly, in our hands none of these mutations resulted in obvious centrosome duplication defects, suggesting a specific defect in Cep135-deficient cells when compared to other MCPH models.

Despite the frequency of centrosomal defects characteristic of the perturbed neural development observed in MCPH patients, it remains to be resolved, however, why brain size in particular is so vulnerable to these defects (Jayaraman et al., 2018, Saade et al., 2018). Our observations in mid-gestation embryos suggest that the abnormal divisions and p53 response to *Cep135*-loss are present in additional tissues such as the embryonic liver. However, p53 induction and apoptosis is only present in the developing cortex at later developmental stages suggesting an adaptive response to lack of Cep135. It thus appears that, compared with other organs, adaptation to centrosome defects is less efficient in the developing neural system, perhaps explaining at least partially the tight correlation between mutation in genes encoding centrosomal proteins and primary microcephaly.

## Materials and Methods

### Generation of mutant mice

Two different CRISPR sgRNAs were designed for generating null alleles as follows: *Cep135*_sg#1: AGTATATTAACATTCGGAAG; *Cep135*_sg#2: CGGAAGAGGTTAGACCAGCT; *Aspm*_sg#1: TGGCGACAAAACGGGATTGA; *Aspm*_sg#2: GGACACGTAGGTCAGCAAAC; *Cdk5rap2*_sg#1 GTCCTTCATGTTCCGGGCTC; *Cdk5rap2*_sg#2: TTCATGTTCCGGGCTCTGGT. Ribonucleoprotein complexes were assembled by incubating Cas9 protein (LabOmics) (100 ng/ml) and 50 ng/ml of each sgRNA in microinjection buffer (10 mM Tris pH 7.4, 0.1 mM EDTA) for 15 minutes at room temperature. The ribonucleoprotein complexes were injected in the cytoplasm of zygotes obtained from matings of B6.CBA males and females at 0.5 days of gestation, using protocols previously described (Henao-Mejia, Williams et al., 2016). 45 zygotes were injected and 28 were transferred to CD1 pseudopregnant females, from which 9 pups were born. Genotyping was performed by PCR and sanger sequencing. Genotyping primers are available upon request. *Trp53* knockout mice were obtained from Jackson Laboratories (B6.129S2-Trp53tm1Tyj/J; JAX stock #002101). Mice were maintained on a C57BL/6J genetic background and were housed in a pathogen-free animal facility at the CNIO following the animal care standards of the institution. The animals were observed on a daily basis, and sick mice were humanely euthanized in accordance with the Guidelines for Humane End-points for Animals Used in Biomedical Research (Directive 2010/63/EU of the European Parliament and Council and the Recommendation 2007/526/CE of the European Commission). For histological examination, samples were fixed in a solution of 4% of paraformaldehyde, embedded in paraffin, and cut into 2.5-μm sections. The sections were then stained with H&E or Cresyl violet (for Nissl staining). All animal protocols were approved by the committee for animal care and research of the Instituto de Salud Carlos III/Comunidad de Madrid (Madrid, Spain).

### Molecular imaging

Computerized axial tomography scanning (CT scan) was performed on anesthetized P30 mice [3% of Isoflurane (Isoba Vet)] using an eXplore ista PET-CTscan (GE Healthcare) with the following parameters: 200mA, 35 kV, 160m, 16 shots and 360 projections. MMWKS (GE Healthcare) and MicroView software were used to analyze the resulting images.

### Immunohistochemistry

Tissues were fixed in 10% buffered formalin (Sigma) and embedded in paraffin for routine histological analysis. Immunohistochemistry was performed on 2μm paraffin sections by using an automated protocol developed for the DISCOVERYXT-automated slide-staining system (Ventana Medical Systems Inc.). All steps were performed in this staining platform by using validated reagents, including deparaffinization, acidic antigen retrieval, and antibody incubation and detection. Primary antibodies used for this purpose are listed in Supplementary Table 10. Appropriate biotinylated secondary antibodies were used to detect the primary antibodies before mentioned, followed by incubation with streptavidin–horseradish peroxidase and diaminobenzidine system. Full slides were digitalized with a Zeiss AxioScan Z1, and analyzed by using the ZEISS Zen 2.3 Imaging Software (Zeiss).

### Immunofluorescence

Immunohistofluorescence was performed in mouse embryos or newborns tissues. Briefly, mouse embryos or newborns (P0) were decapitated and brains dissected out (in the case of P0 newborns). Embryonic heads or P0 brains were fixed in 4% paraformaldehyde (electron microscopy sciences, #50-980-487) for 3h at room temperature and then overnight at 4ºC. Next, tissues were rinsed in PBS (sigma) three times, immersed in 30% sucrose (sigma) for 48h at 4ºC, embedded in Tissue Tek OCT compound (Sakura) and cryosectioned onto SuperFrost Plus slides in a Leica cryostat in 12-14 μm thick cryosections. For immunostaining, cryosections were rehydrated, and boiled in a microwave oven for epitope retrieval (if indicated) in sodium citrate buffer (10 mM [pH 6]) for 1 min maximum. Cryosections were equilibrated in PBS, permeabilized with Triton X-100 0.5% for 5 min, washed, and incubated with blocking solution (4% normal goat serum [Vector Laboratories, S-2000] in PBS-T [PBS with Triton X-100 0.1%]) for 1 h at room temperature. Primary antibodies listed in supplementary table 1 were applied when stated in the indicated concentrations and incubated overnight at 4ºC in a humid chamber. Appropriate secondary antibodies (Alexa Fluor dyes from Molecular Probes) were used at a concentration of 1:500 and incubated RT during 3 hours, followed by washing steps, a labelling step with phalloidin-alexa Fluor-647 (to stain F-actin) or 4′,6-diamidino-2-phenylindole (DAPI, to stain nuclei) when indicated. Slides were mounted using Prolong Gold antifade mounting medium or Fluoromount mounting medium, and visualized in a Leica TCS SP5 confocal laser microscope using a HCX PLAN APO CS 63x/1.4 Oil Immersion, HCX PLAN APO CS 40×/1.4 Oil Immersion or HCX PLAN APO CS 20×/1.4 objective.

For immunofluorescence of cells in culture, cells were grown onto glass coverslips and fixed in PFA 4% for 15 minutes RT or ice-cold methanol for 5 min at −20ºC (when indicated) and then washed in PBS. For neurosphere immunostaining, incipient neurospheres generated during 2 days *in vitro* by clonal dilution of primary NPs. Neurospheres were placed onto matrigel-coated glass coverslips (Corning) for 5 min and fixed immediately with PFA 4%. Further details about the antibodies used in this study are included in Supplementary Table 10.

### Transmission electron microscopy

Cells were grown on P6MW dishes (corning) and fixed with glutaraldehyde 4% (Sigma, # G5882). Fixed cells were washed 3x in 0.1M cacodylate buffer, then post-fixed in 1% osmium tetroxide/1.5% potassium ferrocyanide for 30min. Cells were then washed 3x and incubated in 1% aqueous uranyl acetate for 30 min, washed, and subsequently dehydrated in grades of alcohol (5 min each; 50%, 70%, 95%, 2x 100%). Cells were embedded in TAAB Epon (Marivac Canada Inc.) and polymerized at 60°C for 72 hrs. After polymerization, 1mm squares of the embedded monolayer were glued onto an empty Epon block for sectioning.

For tissue electron microscopy, E14.5 embryonic brains were dissected and fixed immediately in 2% glutaraldehyde (Sigma, #G5882) 4% paraformaldehyde (electron microscopy sciences, #50-980-487) in 0.4M HEPES buffer for 2h RT. Entire embryonic brains were then postfixed with 1% osmium tetroxide (OsO4)/1.5% potassium ferrocyanide (KFeCN^6^) for 1 hour. Then brains were washed in 0.4M HEPES 3x and incubated in 1% aqueous uranyl acetate for 1 hour, followed by 2 washes in 0.4M HEPES and subsequent dehydration in grades of alcohol (10 min each; 50%, 70%, 90%, 2x 10 min 100%). The samples were then infiltrated for 30 min in a 1:1 mixture of propylene oxide and TAAB Epon (Marivac Canada Inc). The samples were embedded in drops of TAAB Epon (Electron Microscopy Sciences) and polymerized in molds at 60°C for at least 72 hours. Ultrathin sections of 40 nm were cut on a Reichert Ultracut-S microtome, placed onto copper TEM grids, stained with uranyl acetate and examined in a Tecnai G2 spirit transmission electron microscope equipped with a lantane hexaboride (LaB6) filament and a TemCam-F416 (4k x 4k) camera with CMOS technology.

### Cell culture and time-lapse microscopy

Primary mouse embryonic fibroblasts (MEFs) were isolated from E14.5 mouse embryos, dissociated by trypsinization, and cultured in Dulbecco’s Modified Eagle’s Medium (DMEM) with high glucose (Sigma, #D6429) supplemented with 10% fetal bovine serum (FBS, Sigma), glutamax supplement (Gibco #35050038) and 0.1% gentamycin. All experiments were performed with primary cultures of passage 0 (P0).

For time-lapse microscopy, MEFs were seeded in IBIDI 8 well µ-slide chambers (IBIDI #80826) at a medium confluency and nuclei stained for 30 min with SiR-DNA kit (Spirochrome) following manufacturer’s instructions. Micrographs in time frames of 10 min were acquired during 24 h in a Deltavision RT imaging system (Olympus IX70/71, Applied Precision) equipped with a Plan apochromatic 20X/1.42 N.A. objective, and maintained at 37°C in a humidified CO^2^ chamber. The resulting videos were processed and analyzed with ImageJ software.

Primary neural progenitors (NPs) were obtained from E14.5 embryos. Briefly, E14.5 embryonic brains were dissected under aseptic conditions; neocortices from dorsal telencephalon were separated and digested with papain (Worthington #LS00310). Papain was activated in an activation solution consisting of 5.5 mM L-cysteine (Sigma, #C8277) and 1.1 mM EDTA (Sigma, #E6511) in EBSS (Gibco #24010-043) following manufacturer’s instructions, filtered through a 0.22 µM filter and used at a concentration of 12 U/ml (1ml/two hemicortices) together with DNAse I (Gibco) for 30 min at 37ºC. NPs obtained were pelleted in DMEM/Hams F-12 (Gibco) and activation solution removed completely. Primary NPs were cultured in suspension in a medium containing DMEM/Hams F-12 (Gibco #11320033), 5 mM hepes, 1 mM sodium pyruvate, 2 mM L-glutamine, N2 supplement (Gibco #17502048), B27 supplement (Gibco #17504044), 0.7 U/ml heparin sodium salt (Sigma #H3149), 20 ng/ml mouse EGF (Gibco #PMG8044), 20 ng/ml mouse FGF2 (Gibco #PMG0034) and penicillin/streptomycin (Gibco #15140148). When needed, neurospheres were dissociated with accutase for 5 min at 37ºC prior to experiment plating.

### Limiting Dilution Assays

Limited dilution assays were performed as previously described (Hu & Smyth, 2009). Briefly, NP cultures were plated at density of 100 cells/ml and incubated for 48 h. Neurospheres formed were dissociated and plated in NPs medium in 96-well plates at different cellular densities (100, 50, 20 and 10 cells per well, respectively). One week later, neurosphere formation was assessed and wells in which there was at least one neurosphere were considered positive. Data in the corresponding representations indicates the fraction of cells with ability to generate cultures with new neurospheres. Graphs were obtained using the ELDA software (Hu & Smyth, 2009) that processes the data obtained in each experimental condition to the limiting dilution model. In these graphs the slopes of the depicted solid lines correspond to the fraction of cells with ability to generate new spheres cultures. A lower slope value indicates a lower fraction of cells with capacity to generate new neurospheres. Dotted lines represent the 95% confidence interval.

### Cell sorting and metaphase spreads

Flow cytometry analysis of DNA content was performed by fixing cells with cold 70% ethanol followed by staining with 10 μg/ml propidium iodide (Sigma). Data acquisition was performed with a Fortessa LSR analyzer (BD Biosciences).

For metaphase spreads, cultured cells were exposed to colcemid (10 µg/ml; #295892 from Roche) for 6 h, pelleted, and individual cell suspension hypotonically swollen in 50 ml of 75 mM KCl at 37ºC for 20-30 minutes. Hypotonic treatment was stopped by gentle addition of 1 ml of Carnoy’s solution (75% pure methanol, 25% glacial acetic acid); cells were then spun down and fixed twice with Carnoy’s solution for 20-30 min at room temperature. After fixation, cells were dropped from a 1-meter height onto clean, prewarmed glass slides. Cells were dried overnight at 70ºC and stained with giemsa (Sigma) afterwards, following standard procedures. Images were acquired with a Leica D3000 microscope and a 60X Plan Apo N (numerical aperture, 1.42) objective. Chromosomes from 100 cells per genotype were counted using ImageJ software.

### 3D immunofluorescence of E14.5 embryos (FLASH)

Whole E14.5 embryos were processed for FLASH (Messal, Alt et al., 2019) by replacing Borate-SDS with a solution of 3 [dimethyl(tetradecyl)azaniumyl]propane-1-sulfonate detergent (80 g/L), Boric acid (200 mM) and Urea (250 g/L)(Tedeschi, Almagro et al., 2020). Immunofluorescence was performed as stated above using the antibodies indicated in Supplementary Table 10. Images were acquired with a light sheet microscope (LAvision Ultramicroscope II). SPIM images 3D reconstruction and surfaces rendering for positive staining was performed with Imaris v9 software (Bitplane).

### RNAseq analysis

Total RNA derived from E11.5 and E14.5 embryonic neocortices and cultured neurospheres was isolated using mirVana kit and sample RNA Integrity was assayed on an Agilent 2100 Bioanalyzer. Sequencing libraries were prepared with the “QuantSeq 3’ mRNA-Seq Library Prep Kit (FWD) for Illumina” (Lexogen, Cat. No. 015) by following manufacturer instructions. Library generation is initiated by reverse transcription with oligodT priming, and a second strand synthesis is performed from random primers by a DNA polymerase. Primers from both steps contain Illumina-compatible sequences. cDNA libraries were purified, applied to an Illumina flow cell for cluster generation and sequenced on an Illumina instrument (see below) by following manufacturer’s protocols. Read adapters and polyA tails were removed with bbduk.sh, following the Lexogen recommendations. Processed reads were analysed with the nextpresso pipeline {Graña, 2018 #193}, as follows: Sequencing quality was checked with FastQC v0.11.7 (http://www.bioinformatics.babraham.ac.uk/projects/fastqc/). Reads were aligned to the mouse reference genome (GRCm38) with TopHat-2.0.10 using Bowtie 1.0.0 and Samtools 0.1.19 (--library-type fr-secondstrand in TopHat), allowing three mismatches and twenty multihits. Read counts were obtained with HTSeq-count v0.6.1 (--stranded=yes), using the mouse gene annotation from GENCODE (gencode.vM20.GRCm38.Ensembl95). Differential expression was performed with DESeq2, using a 0.05 FDR. Enrichr (Kuleshov, Jones et al., 2016) and DAVID (Huang da, Sherman et al., 2009) were used for gene set enrichment analysis of differentially expressed genes. GSEAPreranked were used to perform gene set enrichment analysis for several gene signatures on a pre-ranked gene list, setting 1000 gene set permutations. Only those gene sets with significant enrichment levels (FDR q-value < 0.25) were considered. Access to RNA-seq data is provided from the Gene Expression Omnibus, under the ID GSE14749.

### Statistical analyses

Statistics was performed using Prism software (GraphPad Software) or Microsoft Excel. Unless stated otherwise, all statistical tests of comparative data were done using a Mann-Whitney U test or an unpaired, 2-sided Student’s t-test with Welch’s correction when appropriate. Data are expressed as the mean of at least 3 independent experiments ± SEM, with a *P* value of less than 0.05 considered statistically significant.

## Acknowledgements

We thank Pilar Redondo and Carmen García Martín (CNIO) for technical assistance and acquisition of TEM micrographs, and the Mouse Genome Editing Unit (CNIO) for zygote microinjections for the generation of the CRISPR-edited mutant mice. JGM and DMA received predoctoral contracts from the Ministry of Education of Spain (FPI grants). AWC was supported by Fondation pour la Recherche Médicale and European Molecular Biology Organization (EMBO). JGJ received a short fellowship from EMBO. This work was supported by a grant from the European Commission Seventh Framework Programme (ERA-NET NEURON8-Full-815-094), grants from AEI-MICIU/FEDER (RTI2018-095582-B-I00 and RED2018-102723-T), Worldwide Cancer Research and the AECC (WCR20-155), and the iLUNG programme from the Comunidad de Madrid (B2017/BMD-3884). CNIO is a Severo Ochoa Center of Excellence (MINECO award SEV-2015-0510).

## Author Contributions

JG.-M. performed most of the experiments. A.W.C, D.M.-A, L.R.L.-S., J.G.J. and A.P. participated in the analysis of cellular phenotypes and analysis of embryo abnormalities. J.A. and A.B. performed the whole-mount immunofluorescence studies. D.M. and J.B. helped with fluorescent and electron microscopy, respectively. O.G.-C. helped with the bioinformatics analysis. S.O. contributed to the injection of mouse embryos. M.M. designed the project and J.G.-M. and M.M. wrote the manuscript. All the authors contributed to the analysis of the data.

## References

Barbelanne M, Tsang WY (2014) Molecular and cellular basis of autosomal recessive primary microcephaly. Biomed Res Int 2014: 547986

Bazzi H, Anderson KV (2014) Acentriolar mitosis activates a p53-dependent apoptosis pathway in the mouse embryo. Proc Natl Acad Sci U S A 111: E1491–500

Bettencourt-Dias M, Glover DM (2007) Centrosome biogenesis and function: centrosomics brings new understanding. Nat Rev Mol Cell Biol 8: 451–63

Bettencourt-Dias M, Rodrigues-Martins A, Carpenter L, Riparbelli M, Lehmann L, Gatt MK, Carmo N, Balloux F, Callaini G, Glover DM (2005) SAK/PLK4 is required for centriole duplication and flagella development. Curr Biol 15: 2199–207

Carvalho-Santos Z, Machado P, Alvarez-Martins I, Gouveia SM, Jana SC, Duarte P, Amado T, Branco P, Freitas MC, Silva ST, Antony C, Bandeiras TM, Bettencourt-Dias M (2012) BLD10/CEP135 is a microtubule-associated protein that controls the formation of the flagellum central microtubule pair. Dev Cell 23: 412–24

Carvalho-Santos Z, Machado P, Branco P, Tavares-Cadete F, Rodrigues-Martins A, Pereira-Leal JB, Bettencourt-Dias M (2010) Stepwise evolution of the centriole-assembly pathway. J Cell Sci 123: 1414–26

Dobbelaere J, Josue F, Suijkerbuijk S, Baum B, Tapon N, Raff J (2008) A genome-wide RNAi screen to dissect centriole duplication and centrosome maturation in Drosophila. PLoS Biol 6: e224

Farooq M, Fatima A, Mang Y, Hansen L, Kjaer KW, Baig SM, Larsen LA, Tommerup N (2016) A novel splice site mutation in CEP135 is associated with primary microcephaly in a Pakistani family. J Hum Genet 61: 271–3

Fish JL, Kosodo Y, Enard W, Paabo S, Huttner WB (2006) Aspm specifically maintains symmetric proliferative divisions of neuroepithelial cells. Proc Natl Acad Sci U S A 103: 10438–10443

Gai M, Bianchi FT, Vagnoni C, Verni F, Bonaccorsi S, Pasquero S, Berto GE, Sgro F, Chiotto AM, Annaratone L, Sapino A, Bergo A, Landsberger N, Bond J, Huttner WB, Di Cunto F (2016) ASPM and CITK regulate spindle orientation by affecting the dynamics of astral microtubules. EMBO Rep 17: 1396–1409

Gonczy P (2012) Towards a molecular architecture of centriole assembly. Nat Rev Mol Cell Biol 13: 425–35

Habedanck R, Stierhof YD, Wilkinson CJ, Nigg EA (2005) The Polo kinase Plk4 functions in centriole duplication. Nat Cell Biol 7: 1140–6

Henao-Mejia J, Williams A, Rongvaux A, Stein J, Hughes C, Flavell RA (2016) Generation of Genetically Modified Mice Using the CRISPR-Cas9 Genome-Editing System. Cold Spring Harb Protoc 2016: pdb prot090704

Hilbert M, Noga A, Frey D, Hamel V, Guichard P, Kraatz SH, Pfreundschuh M, Hosner S, Fluckiger I, Jaussi R, Wieser MM, Thieltges KM, Deupi X, Muller DJ, Kammerer RA, Gonczy P, Hirono M, Steinmetz MO (2016) SAS-6 engineering reveals interdependence between cartwheel and microtubules in determining centriole architecture. Nat Cell Biol 18: 393–403

Hu Y, Smyth GK (2009) ELDA: extreme limiting dilution analysis for comparing depleted and enriched populations in stem cell and other assays. J Immunol Methods 347: 70–8

Huang da W, Sherman BT, Lempicki RA (2009) Systematic and integrative analysis of large gene lists using DAVID bioinformatics resources. Nat Protoc 4: 44–57

Hung LY, Tang CJ, Tang TK (2000) Protein 4.1 R-135 interacts with a novel centrosomal protein (CPAP) which is associated with the gamma-tubulin complex. Mol Cell Biol 20: 7813–25

Hussain MS, Baig SM, Neumann S, Nurnberg G, Farooq M, Ahmad I, Alef T, Hennies HC, Technau M, Altmuller J, Frommolt P, Thiele H, Noegel AA, Nurnberg P (2012) A truncating mutation of CEP135 causes primary microcephaly and disturbed centrosomal function. Am J Hum Genet 90: 871–8

Inanc B, Putz M, Lalor P, Dockery P, Kuriyama R, Gergely F, Morrison CG (2013) Abnormal centrosomal structure and duplication in Cep135-deficient vertebrate cells. Mol Biol Cell 24: 2645–54

Insolera R, Bazzi H, Shao W, Anderson KV, Shi SH (2014) Cortical neurogenesis in the absence of centrioles. Nat Neurosci 17: 1528–35

Jayaraman D, Bae BI, Walsh CA (2018) The Genetics of Primary Microcephaly. Annu Rev Genomics Hum Genet 19: 177–200

Jayaraman D, Kodani A, Gonzalez DM, Mancias JD, Mochida GH, Vagnoni C, Johnson J, Krogan N, Harper JW, Reiter JF, Yu TW, Bae BI, Walsh CA (2016) Microcephaly Proteins Wdr62 and Aspm Define a Mother Centriole Complex Regulating Centriole Biogenesis, Apical Complex, and Cell Fate. Neuron 92: 813–828

Kuleshov MV, Jones MR, Rouillard AD, Fernandez NF, Duan Q, Wang Z, Koplev S, Jenkins SL, Jagodnik KM, Lachmann A, McDermott MG, Monteiro CD, Gundersen GW, Ma’ayan A (2016) Enrichr: a comprehensive gene set enrichment analysis web server 2016 update. Nucleic Acids Res 44: W90–7

Leidel S, Delattre M, Cerutti L, Baumer K, Gonczy P (2005) SAS-6 defines a protein family required for centrosome duplication in C. elegans and in human cells. Nat Cell Biol 7: 115–25

Lin YC, Chang CW, Hsu WB, Tang CJ, Lin YN, Chou EJ, Wu CT, Tang TK (2013) Human microcephaly protein CEP135 binds to hSAS-6 and CPAP, and is required for centriole assembly. EMBO J 32: 1141–54

Little JN, Dwyer ND (2019) p53 deletion rescues lethal microcephaly in a mouse model with neural stem cell abscission defects. Hum Mol Genet 28: 434–447

Lizarraga SB, Margossian SP, Harris MH, Campagna DR, Han AP, Blevins S, Mudbhary R, Barker JE, Walsh CA, Fleming MD (2010) Cdk5rap2 regulates centrosome function and chromosome segregation in neuronal progenitors. Development 137: 1907–17

Marjanovic M, Sanchez-Huertas C, Terre B, Gomez R, Scheel JF, Pacheco S, Knobel PA, Martinez-Marchal A, Aivio S, Palenzuela L, Wolfrum U, McKinnon PJ, Suja JA, Roig I, Costanzo V, Luders J, Stracker TH (2015) CEP63 deficiency promotes p53-dependent microcephaly and reveals a role for the centrosome in meiotic recombination. Nat Commun 6: 7676

McKinley KL, Cheeseman IM (2017) Large-Scale Analysis of CRISPR/Cas9 Cell-Cycle Knockouts Reveals the Diversity of p53-Dependent Responses to Cell-Cycle Defects. Dev Cell 40: 405–420 e2

Megraw TL, Sharkey JT, Nowakowski RS (2011) Cdk5rap2 exposes the centrosomal root of microcephaly syndromes. Trends Cell Biol 21: 470–80

Messal HA, Alt S, Ferreira RMM, Gribben C, Wang VM, Cotoi CG, Salbreux G, Behrens A (2019) Tissue curvature and apicobasal mechanical tension imbalance instruct cancer morphogenesis. Nature 566: 126–130

Mottier-Pavie V, Megraw TL (2009) Drosophila bld10 is a centriolar protein that regulates centriole, basal body, and motile cilium assembly. Mol Biol Cell 20: 2605–14

Ohta T, Essner R, Ryu JH, Palazzo RE, Uetake Y, Kuriyama R (2002) Characterization of Cep135, a novel coiled-coil centrosomal protein involved in microtubule organization in mammalian cells. J Cell Biol 156: 87–99

Pulvers JN, Bryk J, Fish JL, Wilsch-Brauninger M, Arai Y, Schreier D, Naumann R, Helppi J, Habermann B, Vogt J, Nitsch R, Toth A, Enard W, Paabo S, Huttner WB (2010) Mutations in mouse Aspm (abnormal spindle-like microcephaly associated) cause not only microcephaly but also major defects in the germline. Proc Natl Acad Sci U S A 107: 16595–600

Roque H, Wainman A, Richens J, Kozyrska K, Franz A, Raff JW (2012) Drosophila Cep135/Bld10 maintains proper centriole structure but is dispensable for cartwheel formation. J Cell Sci 125: 5881–6

Saade M, Blanco-Ameijeiras J, Gonzalez-Gobartt E, Marti E (2018) A centrosomal view of CNS growth. Development 145

Tedeschi A, Almagro J, Renshaw MJ, Messal HA, Behrens A, Petronczki M (2020) Cep55 promotes cytokinesis of neural progenitors but is dispensable for most mammalian cell divisions. Nat Commun 11: 1746

Thornton GK, Woods CG (2009) Primary microcephaly: do all roads lead to Rome? Trends Genet 25: 501–10

