## Supplementary Figures for "Lack of adaptation to centriolar defects leads to p53-independent microcephaly in the absence of Cep135"

by José González-Martínez et al.

### Inventory of Supplementary Information

### Supplementary Figures

Supplementary Figure 1. Generation of *Cep135*-mutant mice and representative pathologies.

Supplementary Figure 2. Transcriptomic profiling of Cep135-mutant developing brains and cultured neurospheres.

Supplementary Figure 3. Cep135 mutant embryos display proliferative defects and p53-dependent cell death in E11.5 embryos.

Supplementary Figure 4. Cep135 mutant MEFs present impaired centriole dynamics, increased duration of mitosis, aberrant mitotic spindles and poliploidy.

Supplementary Figure 5. Cep135 loss does not alter spindle orientation of neural progenitors in the ventricular surface of the developing neocortex.

### Supplementary Tables

Supplementary Table 1. Pathways deregulated in E11.5 Cep135-deficient cortices.

Supplementary Table 2. Pathways deregulated in E14.5 Cep135-deficient cortices.

Supplementary Table 3. Pathways deregulated in E14.5 Cep135-deficient cortices cultured for 12 h.

Supplementary Table 4. Pathways deregulated in E14.5 Cep135-deficient cortices cultured for 24 h.

Supplementary Table 5. Enrichment in transcription factor binding sites in E11.5 Cep135-deficient cortices.

Supplementary Table 6. Enrichment in transcription factor binding sites in E14.5 Cep135-deficient cortices.

Supplementary Table 7. Enrichment in transcription factor binding sites in E14.5 Cep135-deficient cortices cultured for 12 h.

Supplementary Table 8. Enrichment in transcription factor binding sites in E14.5 Cep135-deficient cortices cultured for 24  h.

Supplementary Table 9. Most upregulated genes in Cep135-deficient samples.

Supplementary Table 10. Antibodies used in this work.

Supplementary Figures

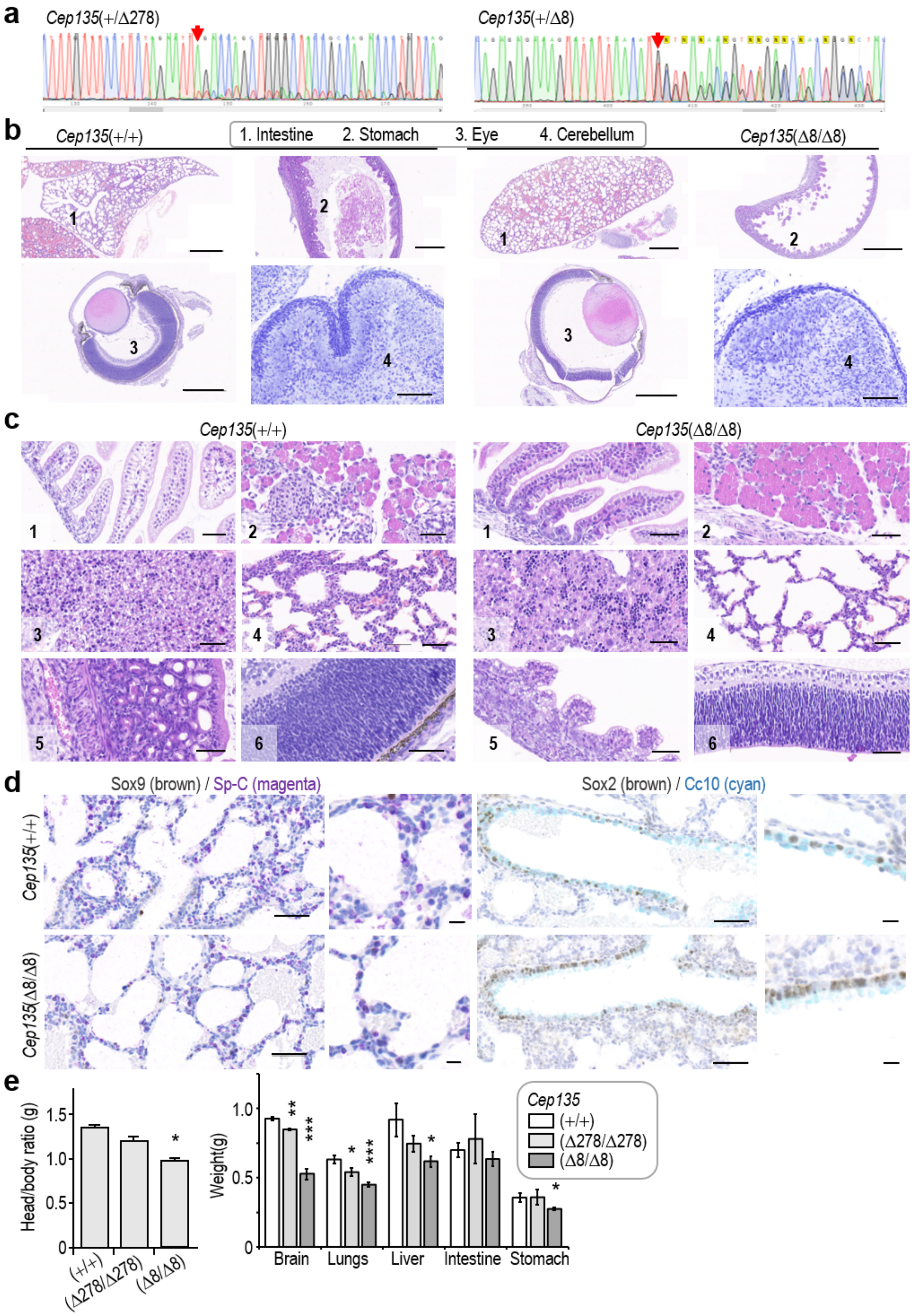


**Supplementary Figure 1. Generation of *Cep135*-mutant mice and representative pathologies.** **a**, Representative sequences corresponding to *Cep135*(+/Δ278) and *Cep135*(+/Δ8) mice. Red arrows indicate indels. **b**, Histological sections stained with hematoxylin and eosin (H&E, 1-3) or Nissl (4) of the indicated organs of P0 *Cep135*(Δ8/Δ8) and control pups. Scale bars: 1 mm. **c**, Higher magnification micrographs of *Cep135*(Δ8/Δ8) and control pups viscerae (1, intestine; 2, pancreas; 3, liver; 4, lung; 5, stomach; 6, retina). Scale bars: 100 µm. **d**, Immunohistochemical staining of *Cep135*(Δ8/Δ8) and control P0 lung sections with the indicated antibodies. Scale bars: 100 µm (micrographs) and 20 µm (higher magnification insets). **e**, Quantification of the brain/body (left) or visceral organs (right) weight ratios in P0 pups of the indicated genotypes. In **e**, data are mean ± SEM from 4 different mouse pups per genotype; *, *P*<0.05; **, *P*<0.01; ***, *P*<0.001 (Student's t-test).

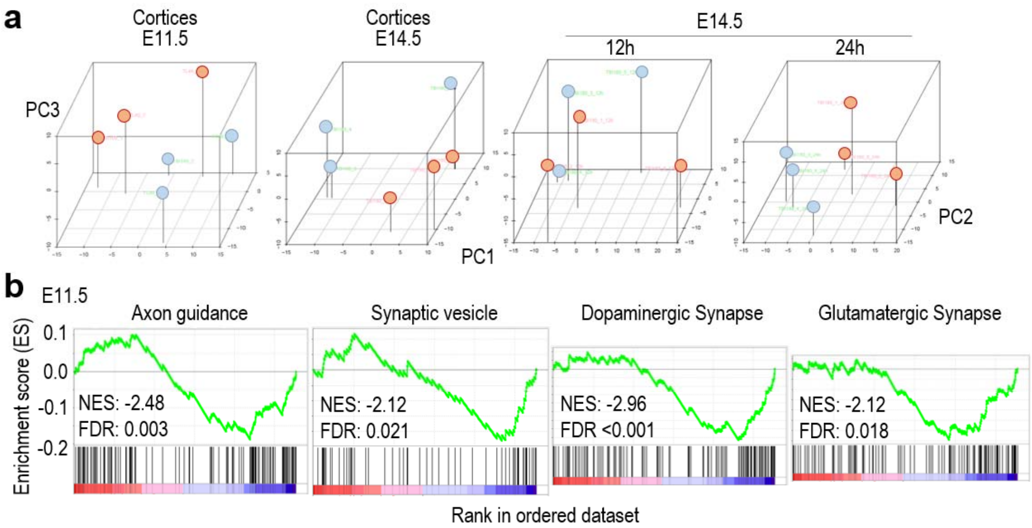


**c**

E14.5 – 0h

| *Deregulated pathways* | |
|---|---|
| *Cluster of Pathways* | *Enrichment Score* |
| Infection and complement | 1.3785 |
| Regulation of transcription | 0.9462 |
| Apoptosis | 0.8141 |
| Protein phosphorylation | 0.3810 |
| G-protein coupled receptors and signal transduction | 0.3144 |

| *Enrichment in TF targets* | |
|---|---|
| Enriched term | *p* value |
| Suz12 Chea | 0.000 |
| AR Chea | 0.001 |
| Rest Chea | 0.003 |
| Tp53 Chea | 0.004 |
| Tp64 Chea | 0.033 |

E14.5 – 12h

| *Deregulated pathways* | |
|---|---|
| *Cluster of Pathways* | *Enrichment Score* |
| Nervous system differ. | 2.1195 |
| cAMP signaling & synapsis | 1.6498 |
| Cell cycle | 1.6477 |
| Ion transport | 1.6048 |
| Synapses signaling | 1.5809 |

| *Enrichment in TF targets* | |
|---|---|
| Enriched term | *p* value |
| Suz12 Chea | 0.000 |
| Smad4 Chea | 0.001 |
| Suz12 ENCODE | 0.009 |
| Rest Chea | 0.010 |
| GATA2 Chea | 0.017 |

**Supplementary Figure 2. Transcriptomic profiling of *Cep135*-mutant developing brains and cultured neurospheres.** **a**, Principal component analysis of the transcriptomic profiles from the indicated *Cep135*(Δ8/Δ8) or *Cep135*(+/+) samples (see Fig. 2 for details). **b**, GSEA analysis of major pathways deregulated in E11.5 *Cep135*(Δ8/Δ8) samples. **c**, Major pathways deregulated and enrichment in transcription factor (TF) targets in E14.5 samples before (0 h) and after being cultured during 12 h to form neurospheres. See Supplementary Tables 1-9 for additional details.

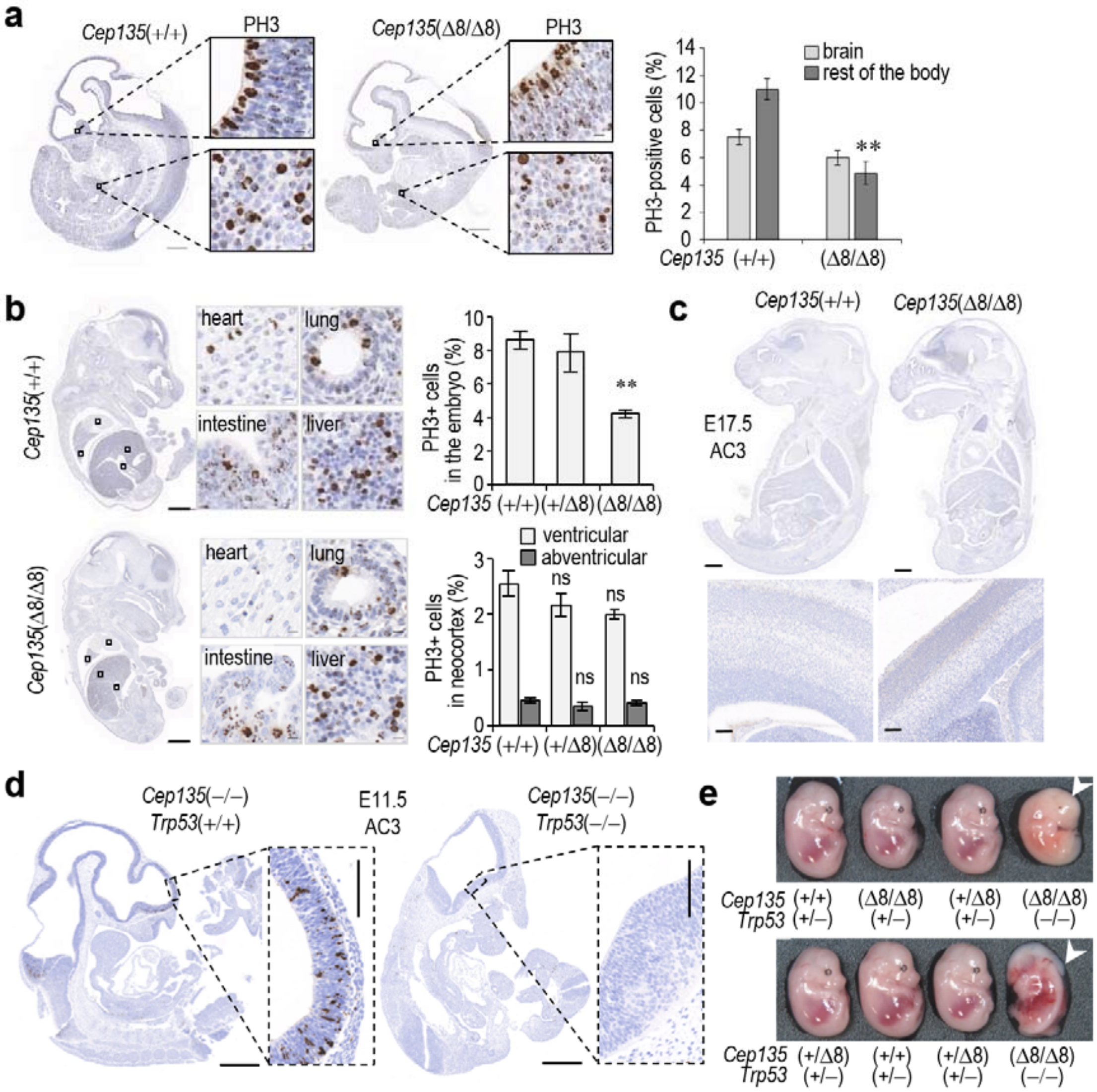


Supplementary Figure 3. *Cep135* mutant embryos display proliferative defects and p53-dependent cell death in E11.5 embryos. **a**, Immunohistochemical staining of phospho-histone H3 Ser10 (PH3) in wild-type and *Cep135*(Δ8/Δ8) E11.5 mouse embryos. The histogram to the right shows the percentage of PH3+ cells in each group. Scale bars: 500 µm (whole embryo sections), 10 µm (insets). **b**, Immunohistochemical staining of PH3 in E14.5 mouse embryos of the indicated genotypes. The histogram to the upper right shows the percentage of PH3+ cells in the embryonic body, and the histogram to the lower right depicts the percentage of ventricular and abventricular mitoses in the dorsal neocortex. Scale bars: 1 mm (whole embryo sections), 10 µm (insets). **c**, Immunohistochemical staining of active caspase 3 (AC3) in E17.5 fetuses of the indicated genotypes. Insets depict higher magnification micrographs of the medial aspect of the dorsal cortex. Scale bars: 1 mm (whole embryo sections), 100 µm (insets). **d**, Immunohistochemical staining of AC3 in E11.5 embryos of the indicated genotypes. Scale bars: 1 mm (whole embryo sections), 100 µm (insets). **e**, Representative bright-field macroscopic images of four E14.5 embryos within two litters resulting from matings between *Cep135*(+/Δ8); *Trp53*(–/–) males and *Cep135*(+/Δ8); *Trp53*(+/–) females. Arrowheads indicate *Cep135*(Δ8/Δ8); *Trp53*(–/–) embryos being reabsorbed. In **a**,**b** data are mean SEM from 3 different embryos per genotype group; **, *P*<0.01; ns: not significant (Student's t-test).

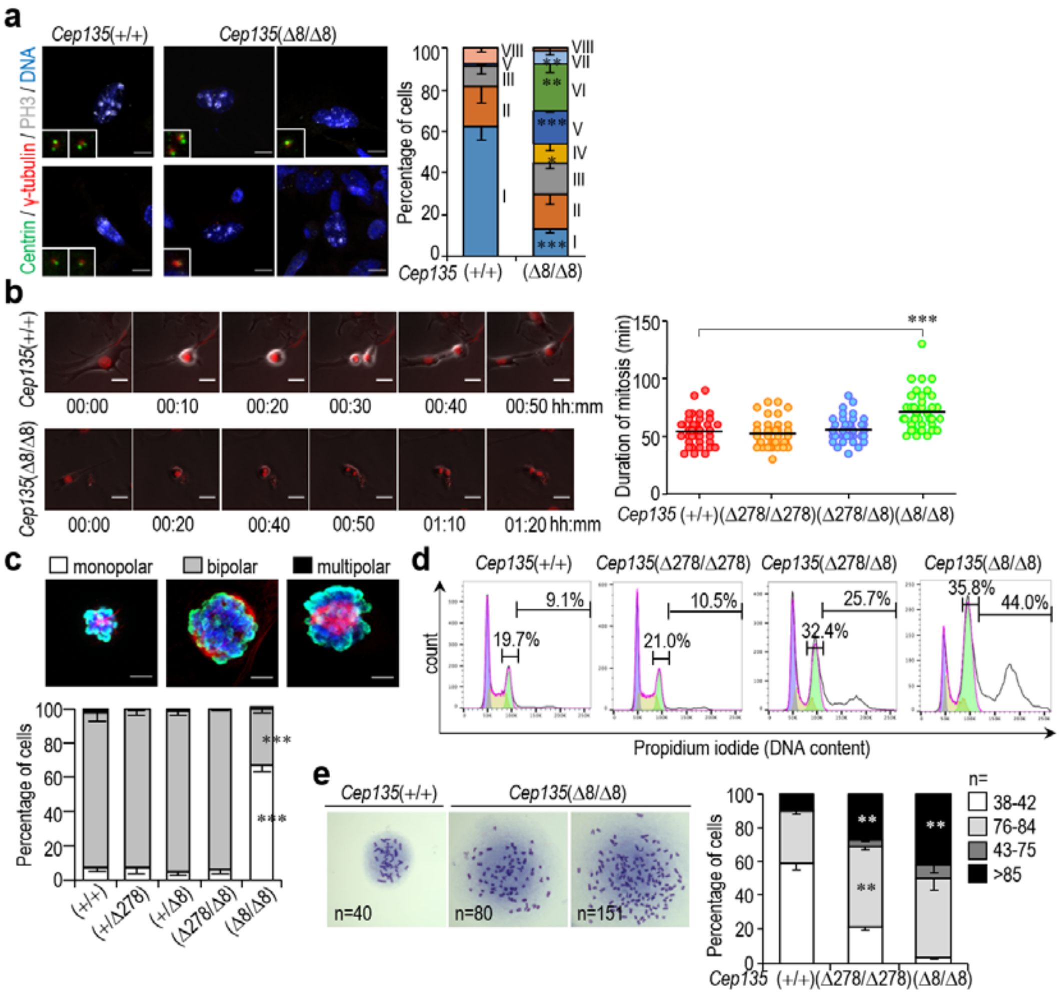


**Supplementary Figure 4.** ***Cep135*** **mutant MEFs present impaired centriole dynamics, increased duration of mitosis, aberrant mitotic spindles and poliploidy.** **a**, Immunostaining for centrin, γ-tubulin and phospho-histone H3 Ser10 (PH3) in E14.5 MEFs of the indicated genotypes. Scale bar: 10 µm. All cells depicted are PH3+. Group I: Two γ-tubulin spots; 2 centrin doublets; II: Two γ-tubulin spots; 2 centrin singlets; III: Two γ-tubulin spots; 1 centrin doublet + 1 centrin singlet; IV: Two γ-tubulin spots; 1 centrin doublet + 1 centrin singlet; V: 1 γ-tubulin spot; 1 centrin doublet; VI: 1 γ-tubulin spot; 1 centrin singlet; VII: acentrosomal; VIII: > 2 γ-tubulin spots. The histogram to the right shows the percentage of cells in each group. **b**, Time-lapse imaging of E14.5 wild-type and *Cep135*(Δ8/Δ8) dividing MEFs. Representative insets depict micrographs for different time points detailed below. The histogram to the right depicts the duration of mitosis (DOM) for the indicated genotypes. Scale bar: 10 µm. **c**, Representative confocal images of monopolar, bipolar and multipolar mitotic spindles in cells stained with the indicated antibodies. The histogram to the bottom depicts the percentage of each mitotic spindle type per genotype. Scale bar: 10 µm. **d**, Cell cycle profiles showing DNA content in E14.5 MEFs of the indicated genotypes. Horizontal lines delimitate the percentage of each population of cells according to their DNA content. **e**, Representative pictures of metaphase spreads of E14.5 *Cep135*(Δ8/Δ8) MEFs and quantification of groups of cells depending on the chromosomic numbers. n indicates number of chromosomes per cell: euploid (2n=40)/tetraploid (4n=80) or near euploid/tetraploid (38-42 and 76-84), or aneuploid (43-75 or >85). In **a**,**c**,**e** data are mean ± SEM; n>150 cells from three independent experiments in **a**, n>50 cells in **b** and **e**, and n>100 cells in **c**. ns, non-significant; **, *P*<0.01; ***, *P*<0.001 (Student's t-test). In **b** horizontal lines represent mean, ***, *P*<0.001 (Unpaired t-test with Welsh correction).

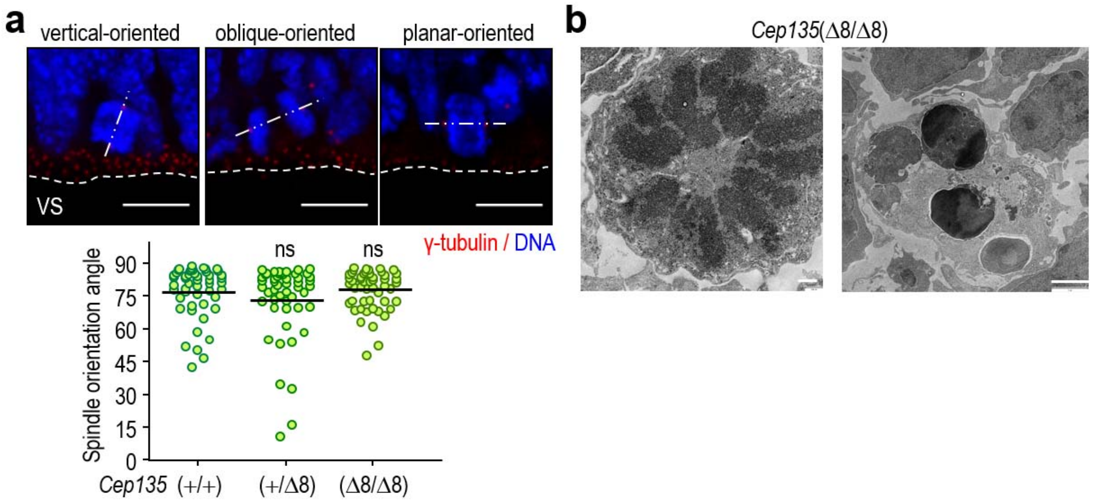


Supplementary Figure 5. *Cep135* loss does not alter spindle orientation of neural progenitors in the ventricular surface of the developing neocortex. a, Representative confocal micrographs of dividing neural progenitors in the ventricular surface (VS) of E14.5 embryos stained for γ-tubulin (red) and DNA (blue). Slashed lines decorating the apical side of γ-tubulin dots represent the VS; dashed and dotted straight lines linking the two centrosomes in each picture represent the approximate orientation of the mitotic spindle. Scale bar: 10 µm. The plot to the bottom depicts the mitotic spindle orientation angle respect the VS; horizontal lines represent mean, ns: non-significant (unpaired t-test with Welsh correction). b, Representative transmission electron microscopy images of monopolar spindles (left) and apoptotic bodies in *Cep135*(Δ8/Δ8) mutant cells in the developing neocortex. Scale bars: 500 nm (left) and 2 µm (right).

### Supplementary Tables

Supplementary Table 1. Pathways deregulated in E11.5 Cep135-deficient cortices (see Excel file).

Supplementary Table 2. Pathways deregulated in E14.5 Cep135-deficient cortices (see Excel file).

Supplementary Table 3. Pathways deregulated in E14.5 Cep135-deficient cortices cultured for 12 h (see Excel file).

Supplementary Table 4. Pathways deregulated in E14.5 Cep135-deficient cortices cultured for 24 h (see Excel file).

Supplementary Table 5. Enrichment in transcription factor binding sites in E11.5 Cep135-deficient cortices (see Excel file).

Supplementary Table 6. Enrichment in transcription factor binding sites in E14.5 Cep135-deficient cortices (see Excel file).

Supplementary Table 7. Enrichment in transcription factor binding sites in E14.5 Cep135-deficient cortices cultured for 12 h (see Excel file).

Supplementary Table 8. Enrichment in transcription factor binding sites in E14.5 Cep135-deficient cortices cultured for 24 h (see Excel file).

Supplementary Table 9. Most upregulated genes in Cep135-deficient samples (see Excel file).

Supplementary Table 10. Antibodies used in this work.

| Antigen | Species | Uses and dilution | Origin |
|---|---|---|---|
| Acetyl-$\alpha$-tubulin | mouse (IgG2b) | IF (1:1000) | Sigma (T7451) |
| $\alpha$-tubulin | mouse (IgG1) | IF (1:1000) | Sigma (T9026) |
| CC10 | goat | IF (1:200) | Santa-Cruz (sc-9772) |
| Cdk5rap2 | rabbit | IF (1:500-1000) | Millipore (06-1398) |
| Cenexin/odf2 | rabbit | IF (1:200) | Abcam (ab43840) |
| Centrin | mouse (IgG2a) | IF (1:500) | Millipore (04-1624) |
| Cep135 | rabbit | IF (1:100) | Abcam (ab75005) |
| Cleaved-caspase 3 | rabbit | IF (1:100), IHC (1:200) | CST (9661S) |
| Cyclin A | rabbit | IF (1:250) | Santa-Cruz (sc-751) |
| $\gamma$-tubulin | mouse (IgG1) | IF (1:1000) | Sigma (T6557) |
| Histone H3 (phospho Ser10) | mouse (IgG1) | IF (1:500) | Millipore (05-806) |
| p53 | mouse (IgG1) | IF (1:200) | CST (2524) |
| PCNA | mouse (IgG2a) | IF (1:200) | Millipore (NA03) |
| Sas6 | mouse (IgG2b) | IF (1:250) | Santa-Cruz (sc-81431) |
| Sox2 | goat | IF, IHC (1:200) | R&D systems (AF2018) |
| SPC | rabbit | IHC (1:100) | Millipore (AB3786) |
| Tbr2 | rabbit | IF (1:250) | Abcam (ab23345) |
| Tuj1 | Mouse (IgG2a) | IF (1:250) | Covance (MMS-435-P) |
| Vimentin (phospho Ser55) | mouse (IgG2b) | IF (1:200) | Abcam (ab23345) |
